# Replication-competent adenoviral platform for in situ production of immunotherapeutic RNA aptamers targeting 4-1BB

**DOI:** 10.64898/2026.03.01.708858

**Authors:** Antonio Carlos Tallon-Cobos, Virginia Laspidea, Iker Ausejo-Mauleon, Daniel de la Nava, Sara Labiano, Marisol Gonzalez-Huarriz, Marta Zalacain, Ana Patiño-Garcia, Helena Villanueva, Juan Fueyo, Candelaria Gomez-Manzano, Ignacio Melero, Fernando Pastor, Marta M Alonso, Marc Garcia-Moure

## Abstract

Viroimmunotherapy leverages oncolytic viruses to induce antitumor immunity and is increasingly explored for solid tumors. Their activity can be enhanced by arming them with immunostimulatory payloads, but most approaches rely on protein-based transgenes that are constrained by viral genome packaging limits. Here, we establish a replication-competent Delta-24-RGD-based platform for localized production of immunotherapeutic RNA aptamers at the tumor site. RNA aptamers provide compact, highly specific ligands that can, in principle, target diverse immune receptors. As a model, we engineered a Delta-24-RGD derivative encoding circular 4-1BB targeting aptamers and show that infected tumor cells sustain aptamer transcription and release, which is associated with a pro-inflammatory remodeling of the tumor microenvironment and measurable antitumor activity in different mouse models with a comparable effect to that achieved with a 4-1BBL-expressing adenovirus used as a benchmark. Overall, this work delivers a proof of concept that replication-competent adenoviruses can serve as in situ factories for extracellularly active RNA aptamers, supporting their development as flexible platforms for localized non-coding cancer immunotherapy.

## INTRODUCTION

The oncolytic virus Delta-24-RGD is a conditionally replicative adenovirus that selectively induces tumor cell death through a dual mechanism: direct oncolysis of infected tumor cells and activation of the antitumor immune response^1^. However, its replication in cancer cells is insufficient to fully overcome the challenges posed by the tumor microenvironment (TME), which limits the spread and persistence of Delta-24-RGD within the tumor mass. As a result, Delta-24-RGD causes only transient infections that are eliminated by the immune system a few weeks after treatment. Cumulative preclinical and clinical evidence highlights the critical role of the immunotherapeutic effects of Delta-24-RGD in promoting durable clinical responses that persist after the virus is cleared^2^.

To further enhance immunostimulatory capacity, research has focused on transforming replicative oncolytic adenoviruses into gene therapy vectors that express immunostimulatory payloads in the TME. The replicative ability of the virus enables a sustained delivery of therapeutic molecules following a single administration. These modified viruses generally surpass the therapeutic effects of the parental virus by increasing immune cell infiltration, proliferation, and immune memory formation^3–7^.

Delta-24-RGD is an attractive gene delivery platform thanks to its intrinsic antitumor properties and favorable safety profile, even in difficult-to-treat tumors like pediatric brainstem cancers^8^. Additionally, it benefits from decades of research into the genetic engineering of human adenovirus serotype 5 (HAdV-C5) vectors, which has yielded highly efficient protocols for modifying its genome and producing the virus. However, a significant constraint of Delta-24-RGD is the restricted carrying capacity of the genome^9^. These size constraints prevent the insertion of large transgenes encoding therapeutically relevant molecules, such as monoclonal antibodies (mAbs) or chimeric constructs.

Agonist and antagonist mAbs are currently the gold standard in immunotherapy owing to their specific, high-affinity binding to virtually any target^10^. However, mAb-based anti-cancer therapies typically require multiple systemic doses, leading to on-target/off-tumor toxicities^11–13^. While intratumoral delivery might reduce some of these toxicities, repeated dosing is not feasible for hard-to-reach tumors^14^. MAbs are ideal candidates for loading into oncolytic viruses to enable sustained intratumoral drug delivery after a single injection, thus minimizing systemic toxicities. However, the size of mAb transgenes must include sequences encoding the heavy chain (50 kDa) and light chain (25 kDa), and their expression cassettes are too large for Delta-24-RGD. Therefore, smaller therapeutic molecules that retain the versatility of mAbs must be explored to effectively vehiculize them within the Delta-24-RGD virus.

Aptamers are small ssDNA or ssRNA oligonucleotides (20-60 nt) with a particular tridimensional structure that allows its specific binding to their target^15^. Aptamers are obtained by the Systematic Evolution of Ligands by Exponential Enrichment method (SELEX) from random oligonucleotide libraries with a high diversity in unique sequences and 3D conformations^16^. Hence, virtually any molecule can be virtually targeted by an aptamer. Because of their versatility and smaller size, aptamers pose an unexplored feasible alternative to mAbs to arm the Delta-24-RGD genome with immunotherapeutic transgenes. Once injected into the tumor, the modified virus will replicate and kill tumor cells while producing and releasing the therapeutic aptamer in situ to potentiate the antitumor immune response.

4-1BB (CD137) is a costimulatory molecule selectively expressed on antigen-primed T lymphocytes, whose function can be modulated with stimulatory ligands such as trimers of its natural ligand or agonist antibodies^17^. Anti-CD137 agonist antibodies in mice bearing transplanted tumors may result in immune-mediated regression of tumors in several instances^18,19^. Multiple agents based on CD137 agonism in the form of monospecific or bispecific antibodies currently are undergoing phase 1 to 3 clinical development^20^.

Here, we present a Delta-24-RGD derivative engineered to express a 4-1BB agonistic RNA aptamer. The new virus serves as a proof of concept to validate the adenovirus Delta-24-RGD as a suitable platform for delivering therapeutic aptamers with immunostimulatory properties to the tumor microenvironment. We demonstrate that the aptamer-expressing virus given intratumorally reduces tumor growth and extends survival in several immunocompetent mouse models in a 4-1BB-dependent manner without compromising viral replication and overall oncolytic properties.

## MATERIALS AND METHODS

A complete list of oligonucleotides (**Table S1**) sequences and antibodies (**Table S2**) can be found in supplemental materials.

### Cell lines and culture conditions

A549 (CCL-185TM), HEK293 (CRL-1573), HCT116 (CCL-247) and K7M2 (CRL-2836) cells were obtained from the American Type Culture Collection (ATCC) and cultured according to the specifications. MDA-BoM-1833, CT26 and 4T1 cell lines were kindly provided by Dr. Fernando Pastor and were cultured with RPMI-1640 Gluta-Max medium supplemented with 10% FBS and 1% antibiotic. 531MII, a primary osteosarcoma cell line, was developed at Clínica Universidad de Navarra were cultured in minimum essential medium supplemented with 10% FBS and 1% antibiotic. Mouse primary lymphocytes were cultured in T-cell media (RPMI, β-mercaptoethanol 1 μM, 1% glutamine, 1% non-essential amino acids, 1% pyruvate, 1 mM HEPES, and 1% P/S, and 10% FBS).

Cell cultures were incubated at 37°C in a humidified atmosphere containing 5 % CO2. All cells were authenticated by Short Tandem Repeat DNA profiling at the CIMA Genomic Core Facility (Pamplona; Spain) and were routinely tested for mycoplasma (Mycoalert; Lonza).

### Animal studies

Tumors were measured along the two perpendicular diameters with a caliper, and calculated the tumor volume with the following formula: Volume (mm3) = D x d^2^ x 0.5, where D is the largest diameter and d is the smaller diameter. Animals were sacrificed when the tumor volume reached 2000 mm^3^ (subcutaneous) or 430 mm^3^ (tibial), or when showing ulceration or symptoms of progressive disease.

### SELEX Protocol

The selection of m41BB-binding aptamers was performed by SELEX. Briefly, a dsDNA oligonucleotide template (Integrated DNA Technologies) containing 40 random nucleotides was transcribed in vitro (AmpliScribe™ T7 High Yield Transcription Kit; Lucigen) and purified by PAGE in denaturing conditions to generate the aptamer library. One nmol of the library was incubated with IgG1a-coupled Sepharose-protein-A beads (Sigma) to counter-select non-specific binders. The remaining unbound molecules in the supernatant were incubated with 10 µg of murine 4-1BB-Fc (R & D systems) coupled Sepharose-protein-A beads. Then, the RNA bound to the 4-1BB protein was pulled down by centrifugation of the Sepharose beads and extracted with phenol/chloroform. The recovered RNA was reverse transcribed (Roche, Ref 10109118001) using the Sel_3 primer and amplified by PCR (Invitrogen) using the primers Sel_5 and Sel_3. The PCR products obtained after column purification were used as the template for the next SELEX round, and the whole process was repeated for six rounds. PCR number of cycles was minimized in each selection round to reduce PCR bias that can lead to unspecific aptamer enrichment. SELEX was performed in a physiological binding buffer—150 mM NaCl, 2 mM CaCl_2_, 20 mM HEPES, and 0.01% BSA—at pH 7.4 at 37°C. For every round of SELEX, 1 nmol of RNA library was incubated with 4-1BB-Fc coated beads for 30 and washed three times with binding buffer. Restriction in the aptamer selection along the SELEX procedure was increased in each round by tripling the binding volume, which resulted in a reduced aptamer library concentration for each new round of selection.

### SELEX sequencing and analysis

The PCR products of the aptamer libraries obtained in rounds 3, 4, 5 and 6 were sequenced using NGS (Ion S5, Thermo Fisher) and subsequently analyzed by two different bioinformatic pipelines based on FASTAptamer^36^ and Aptaguide software to ensure that both analyses converge in the same enriched sequences. FASTAptamer software provides the FASTAcount file that indicates the aptamer ranking enrichment. This file was further analyzed using Clustal Omega to provide all the aptamer families and the dendrograms depicting similarities between the enriched aptamers. RNA aptamer secondary and tridimensional structures were predicted and modeled using RNAstructure^37^ and RNAComposer^38^, respectively. 3D models were visualized using the Mol* 3D Viewer^39^.

### Transcription of 4-1BB RNA aptamers

Monomeric RNA aptamers were transcribed following the same protocol as in the SELEX but replacing the random region in the DNA oligonucleotide template with the specific sequence for each aptamer (Integrated DNA Technologies).

### Binding affinity

Interaction and affinity between RNA aptamers and 4-1BB were assessed by MicroScale Thermophoresis (MST). Briefly, recombinant mouse 4-1BB protein was fluorescently labeled (Protein Labeling Kit RED-NSH 2nd Generation, NanoTemper Technologies). Next, up to 16 serial dilutions of the RNA aptamer (starting from 2.5 µM) and the labeled protein (20 nM) were incubated in Binding Buffer (20 mM HEPES, pH 7.4; 150 mM NaCl; 2 mM CaCl2; 0.01% BSA) + 0.05% Tween-20 at 37 °C for 1 h. Then, the samples were loaded in capillary tubes (Monolith NT.115 Premium Capillary, NanoTemper Technologies) and binding kinetics and K_d_ were measured running a Binding Affinity test (Monolith NT.115, NanoTemper Technologies) at 37 °C.

### Biotinylation of RNA aptamers

Aptamer DNA templates were amplified by PCR (Invitrogen), using the primers Sel5 and Sel3Bio. Sel3Bio is a modified version of the Sel3 primer that adds a short overhang at the 3’ end. The modified templates were transcribed, and the RNA aptamer was purified by PAGE as explained above. Then, 1 nmol of aptamers were incubated with 2 nmol of a 5’ biotinylated ssDNA probe at 85 °C in 100 µL of annealing buffer (NaCl 150 mM, EDTA 10 mM, TRIS-Cl 10mM, pH 7.5), and the temperature was lowered by 2°C every 2 minutes to allow annealing. Then, the biotin-aptamer complexes were concentrated and equilibrated to PBS Ca^2+^Mg^2+^ using a 30 kDa centrifugal filter (Millipore).

### Binding of aptamers to 4-1BB coated beads and lymphocytes

*Beads:* A total of 5 µg of 4-1BB-Fc (R&D Systems, Ref 937-4B-050) or IgG1a (Sigma-Aldrich, Ref I5154) were coupled to 4x10L Dynabeads M-450 Tosylactivated (Invitrogen) following the manufacturer instructions. For each binding assay, 200,000 4-1BB-coated beads were used.

*Lymphocytes:* CD8+ T cells were isolated from the spleens of C57Bl/6 mice (Envigo) using the CD8a+ T Cell Isolation Kit (Ref #130-104-075; Miltenyi Biotec). T cells were either left non-activated (naïve) or activated for 48 hours. Activation was performed by plating 100,000 cells per well in 96-well U-bottom plates pre-coated with 2 µg/mL anti-CD3ε antibody in 200 µL of T-cell culture media supplemented with 2 µg/mL CD28 antibody. For binding assays, 100,000 CD8+ T cells were used.

*Aptamer Labeling and Binding Assay:* 25 pmol of biotinylated aptamers were labeled with PE-labeled streptavidin (BioLegend, Ref 405203). The labeled aptamers were incubated with either 4-1BB-coated beads or CD8+ T cells in a total volume of 100 µ PBS (with Ca² and Mg²) for 30 minutes at 37 °C. After incubation, the samples were washed and analyzed by flow cytometry (CytoFLEX LX) and processed using FlowJo software.

### Plasmid construction, virus generation and production

DNA fragments containing the expression cassettes for dimeric 4-1BB (or a scrambled sequence) were engineered using the U6+27 promoter, Tornado ribozymes, and transcription terminator scaffold sequences were obtained from the pAV-U6+27-Tornado-Broccoli plasmid deposited by the S. Jaffrey lab in Addgene (#124360). Same U6 promoter and terminator sequences were used for the non-Tornado dimeric aptamers. The transgenes were synthesized and cloned into the adenoviral shuttle vector pAB26-RGD (a generous gift from Drs. Fueyo and Gomez-Manzano, MDACC) using ClaI and XbaI restriction sites (GenScript).

Subsequently, EcoRV restriction fragments of the modified plasmids, carrying the expression cassettes, were subcloned into the plasmid pVK-500c-D24 containing the parental adenovirus genome (provided by Drs. Fueyo and Gomez-Manzano, MDACC), previously linearized by SwaI digestion, through homologous recombination in the recombinase-positive BJ5183 bacterial strain (Addgene). The same cloning strategy was followed to generate the adenovirus encoding the 4-1BB aptamer without the Tornado sequences. This process generated the Delta-24-Apt, Delta-24-AptT and Delta-24-ScrT genomes that were finally linearized (PacI) and transfected into A549 cells (Lipofectamine 2000, Invitrogen) for virus amplification and purification by CsCl gradient ultracentrifugation at the Viral Vector Production Unit (UAB, Spain).

### Comparison of 4-1BB_Apt3 expression between Tornado and non-Tornado constructs

HEK293 cultures (500,000 cells) were transfected with pUC plasmids expressing the different aptamers (Lipofectamine 2000), and 48 hours later the cell pellets were harvested, and total RNA was isolated with Trizol (Invitrogen). RNA aptamers and GAPDH were measured by qPCR using the following primers: 4-1BB_Apt3_Dimer Fwd/Rv, Scr_Dimer Fwd/Rv, GAPDH Fwd/Rv.

### RNA expression in virus infected cells

200,000 MDA-BoM-1833 cells were plated in 6-well plates and infected with the corresponding oncolytic virus (MOI 25). Cell pellets were harvested at 16 and 72 hours after infection, total RNA was extracted by Trizol protocol (Invitrogen). 4-1BB Apt3T, GAPDH and Fiber cDNA were quantified by qPCR using the following primers: 4-1BB Apt3T Fwd/Rv, GAPDH Fwd/Rv, Fiber Fwd/Rv.

### Pull-down of 4-1BB_Apt3T aptamer from infected cells

A total of 2 × 10 MDA-BoM-1833 cells were plated in a 100 mm dish and infected with Delta-24-AptT at an MOI of 25. After 72 hours, total RNA was extracted from the cell pellet (TRIzol, Thermo Fisher Scientific, Ref 15596026) and the supernatant (TRIzol LS, Thermo Fisher Scientific, Ref 10296010) following the manufacturer protocol. The RNA was treated with DNase I, refolded by heating at 65°C for 5 minutes and cooling to room temperature, and resuspended in binding buffer containing RNase inhibitor. A 10% fraction of bulk RNA was saved as the input sample, while the remaining RNA was divided into two equal parts and incubated with either 5 µg of 4-1BB-Fc recombinant protein 4-1BB-Fc (R&D Systems, Ref 937-4B-050) or IgG1a (Sigma-Aldrich, Ref I5154) in a final volume of 300 µL of binding buffer at 37 °C for 1 hour under rotation. RNA-protein crosslinking was performed on an ice-cold plate under UV light at 500 kJ/m². Samples were sonicated 20 times for 10 seconds each, with 30-second intervals between pulses. Protein-RNA complexes were precipitated using protein G beads, followed by three washes with binding buffer. The bound fraction was collected and subjected to reverse crosslinking in a buffer containing 2.5 mg/mL proteinase K (Roche, Ref 3115828001), 0.6 % SDS, and 60 mmol/L Tris-HCl (pH 7.4), and the samples were heated at 65°C for 45 minutes. RNA was extracted from both the bound and unbound fractions using TRIzol and analyzed via qRT-PCR to quantify the aptamer 4-1BB_Apt3T using specific forward and reverse primers. For each sample, the Ct value of the aptamer was normalized to GAPDH by calculating 2^-ΔCt. These values were then referenced to the Input sample by dividing each by the 2^-ΔCt of the Input method to determine relative abundance (2^-ΔΔCt).

### T-cell activation with 4-1BB_Apt3T aptamer

CD8+ T cells were isolated from the spleens of naive Balb/c mice using the Miltenyi Mouse CD8+ Isolation Kit. A total of 2x10L CD8+ T cells were seeded in 96-well plates pre-coated with 1 μg/mL anti-CD3ε antibody in T cell culture media. Treatments included 2 μg of purified RNA from MDA-BoM-1833 cells infected with Delta-24-ScrT or Delta-24-AptT viruses at an MOI of 25 for 72 hours, mock-infected controls, 2 μg/mL of 4-1BB agonistic antibody, or an isotype control antibody. After 72 hours, cells were stained with anti-CD25-APC antibody, detected using a CytoFLEX LX instrument (Beckman Coulter), and analyzed with FlowJo software (BD Biosciences).

### Virus replication assays

A549 or MDA-BoM-1833 cells were seeded in 6-well plates at a density of 50,000 cells per well and infected with the corresponding viruses at an MOI of 10. Cells were harvested at 16-, 48-, or 72-hours post-infection, depending on the experiment. For viral genome analysis, cell pellets were collected, washed with PBS, and subjected to total DNA (QIAamp DNA Blood Mini Kit, Ref 51104). Viral and genomic DNA levels were quantified by qPCR using fiber and LDLR primers, respectively. For infectious viral particle analysis, the entire cell-supernatant culture was harvested and subjected to three freeze-thaw cycles. Infectious viral titers (PFU) were determined by incubating HEK293 cultures with serial dilutions of the lysates, followed by detection using the anti-hexon method^40^.

### Viral genomes in subcutaneous tumors

MDA-BoM-1833 tumors (500,000 cells) were subcutaneously engrafted in the flank of NSG mice. When tumor reached 50 mm^3^, 5x10^7^ PFU of Delta-24-AptT were injected in the tumor, and tumors were collected at days 2, 7 and 14 after injection (n = 4-7 per time point). Total tumor DNA was extracted (QIAamp DNA Blood Mini Kit, Ref 51104) and viral load was determined by qPCR (fiber primers) in 50 ng of total DNA.

### Cell viability assays

CT26, 4T1, K7M2, HCT116, MDA-BoM-1833 and 531MII cells were plated in 96-w plates (2,000 cells/well) and treated with increasing doses of the oncolytic viruses ranging from 0.01 to 10,000 PFU/cell in 100 µL of their respective culture media. Cell viability was assessed three days later using the CellTiter 96 Aqueous One Solution Cell Proliferation Assay (G3581, Promega). IC_50_ was determined by non-linear regression (GraphPad Prism).

### Antitumor assessment in vivo

To perform evaluation of antitumor efficacy of the oncolytic viruses in subcutaneous models, 4- to 6-week-old male and female Balb/c mice (Envigo) were inoculated in the right flank with 50 μL of PBS containing 500,000 (CT26) or 50,000 (4T1) cells. When tumor volume reached 50 mm^3^, animals were randomized and treated with an intratumor injection of 5x10^7^ PFU of virus diluted in 10 μL of PBS (day 0). For the orthotopic osteosarcoma model, 500,000 cells (K7M2) were injected in the tibial crest of 6- to 8-week-old mice (Balb/c, Envigo). Then, the animals were randomized and 10^8^ PFU of the corresponding viruses were injected in the same place at days 10 and 18 after tumor injection (days 0 and 8 after treatment, respectively).

### Histological analyses

4T1 tumors were excised when reached endpoint criteria and fixed during 48 h in 4% formalin. Then, samples were incubated in 70% ethanol for 48 h and embedded in paraffin. Formalin-fixed paraffin-embedded (FFPE) tumors samples were sectioned (4 μm) and processed for immunohistochemistry (IHC) against CD3 and CD8 markers at the Morphology Unit. Stained samples were then scanned (Leica Aperio CS2) and positive cells were counted using Fiji platform (ImageJ).

### Immunophenotyping

4T1 tumors were inoculated and treated with the oncolytic viruses as explained above. At day 10 after treatment, the tumors were collected for tumor infiltrating lymphocytes (TILs) isolation. In brief, tumor samples were mechanically and enzymatically disaggregated in RPMI containing 40 μg/mL DNAse I and 1 mg/mL collagenase (Roche) for 20 min at 37°C in constant rotation and homogenized using GentleMACS dissociator (Miltenyi Biotec). Tissue homogenates were filtered through a 40-μm cell strainer to isolate individual cells and centrifuged. Cell pellets were resuspended in 10 mL of 35% Percoll solution (GE Healthcare) and centrifuged to obtain TILs.

TILs were stained with PromoFluor-840 viability dye (PK-PF840-3-01, PromoCell) following the recommended protocol for death cell exclusion in the analyses. Then, cell membrane markers were labelled with Fc block (BD Bioscience, Ref 1293778) and the corresponding conjugated antibody panels (**Table S2**). Finally, the samples were fixed and permeabilized for intracellular staining and analyzed in a CytoFLEX LX device (Beckman Coulter).

### Statistical analysis

Statistical analyses were conducted using GraphPad Prism software. Comparisons between two groups were performed using a two-sided Student’s t-test, while comparisons among multiple experimental groups were analyzed using one-way ANOVA with Tukey’s post-hoc test for pairwise comparisons. For experiments involving two variables (e.g., time and treatment), two-way ANOVA with Tukey’s post-hoc test was used. Non-significant differences were considered for p values higher than 0.05. All statistical analyses were based on, at least, three independent experiments. The specific statistical test and sample size for each experiment are detailed in the corresponding figure legends.

## RESULTS

### Selection of high-affinity and specific m4-1BB targeting aptamers

The presence of modified 2’-fluoropyrimidines in the already published mouse 4-1BB agonist aptamer^21^ precludes its use in adenovirus-infected cells. To overcome this, we selected a new 4-1BB-targeting RNA aptamer by SELEX using unmodified nucleotides, thereby ensuring compatibility with cell-endogenous adenoviral gene expression systems. We constructed a DNA oligonucleotide template containing 40 random nucleotides flanked by constant regions at the 5’ and 3’ ends to achieve this. A T7 RNA polymerase promoter was incorporated at the 5’ end to facilitate in vitro transcription of the RNA aptamer library (**Supplemental Fig S1A**).

This aptamer library underwent six iterative rounds of SELEX against the extracellular domain of m4-1BB, with increasing stringency at each step, resulting in the enrichment of 4-1BB-specific RNA sequences (**Supplemental Fig S1B**). After six rounds of SELEX, 1489 unique sequences were recovered and grouped into 132 distinct families (**Supplemental Fig S1C**). From these, we selected six aptamer candidates (4-1BB_Apt1–6) representing the most frequent sequences from the top six families (**Supplemental Fig S1A and Fig S2**) for further validation.

The affinity of the RNA aptamer candidates for 4-1BB was assessed using micro-scale thermophoresis (MST). Among them, 4-1BB_Apt3 and 4-1BB_Apt5 demonstrated high affinity, with dissociation constants (K_D_) of 290 pM and 550 pM, respectively (**Supplemental Fig S3A**). Notably, 4-1BB_Apt1 and 4-1BB_Apt6 also exhibited strong affinities, both in the low nanomolar range, with K_D_ values below 10 nM. In contrast, 4-1BB_Apt2 had a considerably weaker affinity (51.63 nM), while 4-1BB_Apt4 was the weakest binder, with a K_D_ of nearly 300 nM.

Fluorescently labeled RNA aptamers 4-1BB_Apt3, 4, 5, and 6 were detected in over 90% of m4-1BB-coated beads, with a fold-change in mean fluorescence intensity (MFI) ranging from 2.4 to 3.4 compared to the initial aptamer library (**Supplemental Fig S3B and S3C**) as measured by flow cytometry. 4-1BB_Apt1 displayed reduced binding (77 %, MFI x 1.4), while 4-1BB_Apt2 showed minimal binding.

Next, we determined the capacity of the aptamers to bind activated CD8+ T cells. Again, 4-1BB_Apt2 showed the lowest binding to CD8 cells among all candidates (**Fig 1B and 1C**). 4-1BB_Apt3 and Apt4 were detected in 58% of activated CD8 T cells, approaching saturation levels indicated by the anti-4-1BB antibody (65%) and showing the highest MFI fold change. The binding of 4-1BB_Apt1, Apt5, and Apt6 to activated T cells was considerably lower in the percentage of labeled cells and MFI. None of the aptamers bound naïve CD8 T cells, confirming their specificity for activated lymphocytes expressing 4-1BB. (**Supplemental Fig S3D**). Considering the binding affinity and performance in activated T cells, 4-1BB_Apt3 was selected to generate the oncolytic virus.

**Figure 1.**
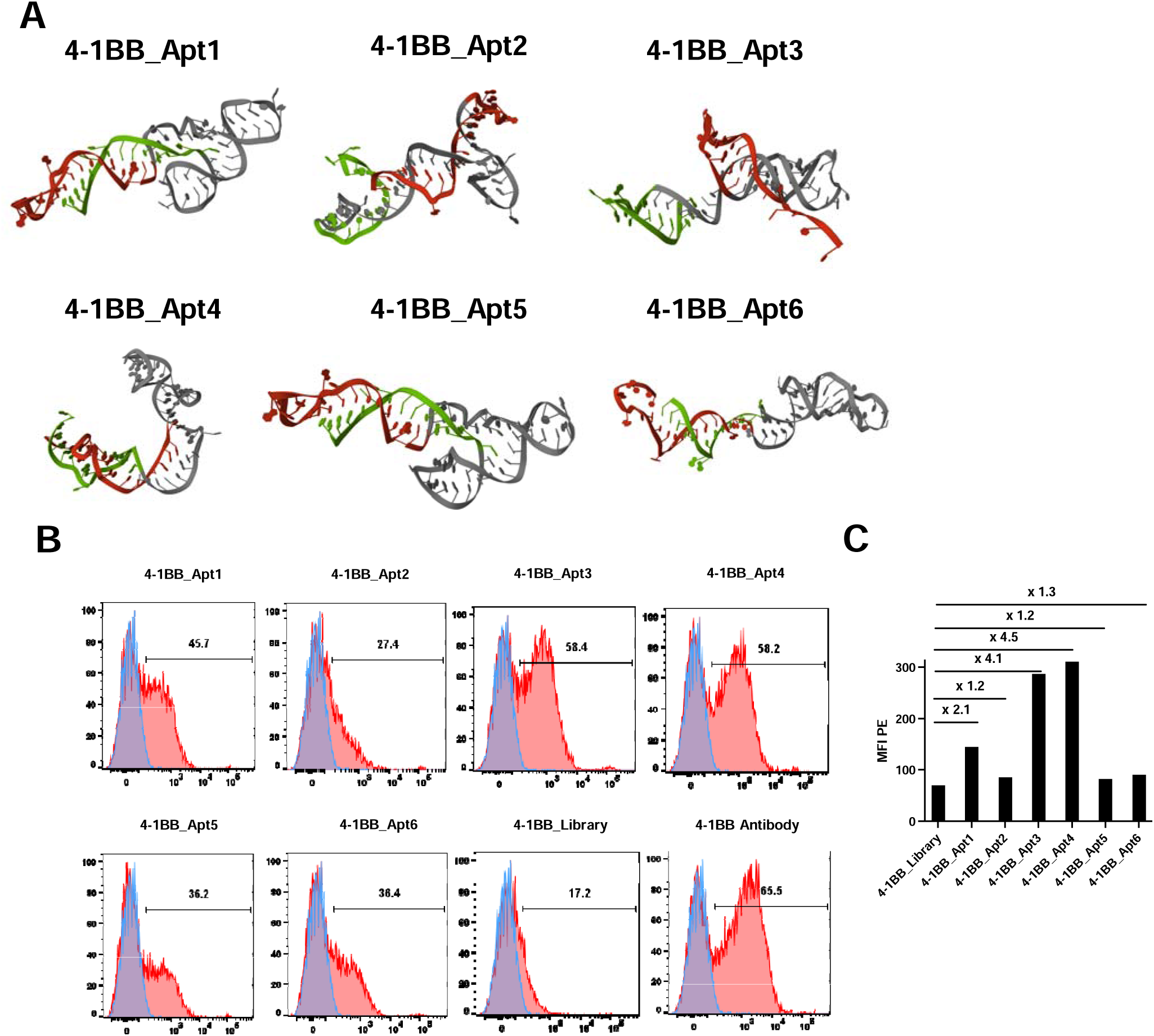
Screening of 4-1BB aptamer candidates. **A)** Predicted tridimensional structure of 4-1BB_Apt1-6. Green and red sequences indicate the 5’ and 3’ constant regions, respectively (See also Supplemental Figure S2). **B)** Flow cytometry histograms showing the binding of PE-fluorescently labeled aptamers 4-1BB_Apt1 to Apt6, 4-1BB_Library and anti-4-1BB antibody (red) or biotin probe (blue) to activated CD8+ T cells. Values indicate the percentage of positive cells. **C)** Median Fluorescence Intensity (MFI) and fold change compared to 4-1BB_Library (randomized initial library used for the aptamer selection). See also Supplemental Figure S3D.

### Design of transgenes expressing 4-1BB aptamer

4-1BB_Apt3 was subjected to two additional modifications to improve its therapeutic effect once produced from virus-infected cells (**Supplemental Fig S4A**). The 4-1BB signaling cascade is triggered by the multimerization of the receptor^21–24^. Thus, first, we engineered a dimeric 4-1BB_Apt3 aptamer by placing a self-complementary RNA linker between two 4-1BB_Apt3 monomers to enable multimerization. Second, we implemented the Tornado system, which incorporates self-complementary autocatalytic ribozymes flanking the aptamer to enable their covalent circularization^25^. This design prevents aptamer degradation by RNA exonucleases, thus extending its half-life. RNA structure simulation verified that the 4-1BB_Apt3 sequences maintain their original folding (**Supplemental Fig S4B**). After confirming that the Tornado element enhances the amount of aptamer (**Supplemental Fig S4C**) and observing no significant differences in virus replication between adenoviruses with and without the Tornado element (**Supplemental Fig S4D**), we proceeded with the 4-1BB_Apt3T-expressing virus Delta-24-AptT) as our candidate of choice.

### Delta-24-AptT infects tumor cells and releases functional 4-1BB Apt3T dimers while maintaining the oncolytic properties

In Delta-24-AptT-infected cells, 4-1BB Apt3T expression showed a significant 6250-fold rise at 72 h compared to 16 h (p = 0.002) (**Fig 2A**), correlating with the temporal increase of the viral late gene fiber (**Fig 2B**). Consistent fiber expression in Delta-24-RGD, Delta-24-ScrT, and Delta-24-AptT indicates that aptamer expression cassettes do not interfere with viral dynamics (**Fig 2C**).

**Figure 2.**
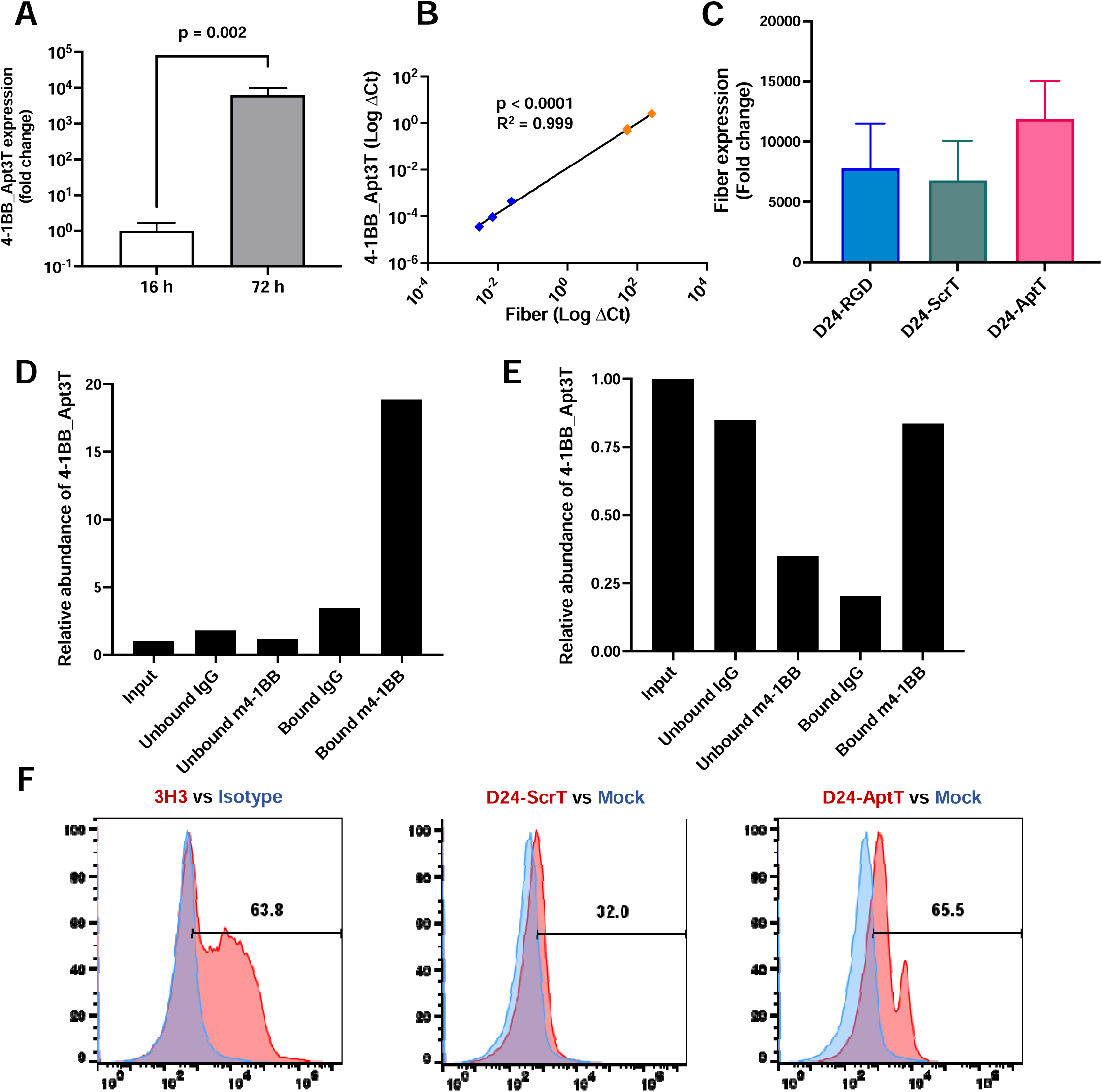
Expression of functional 4-1BB aptamers in Delta-24-Apt3T infected cells. **A)** Increase in 4-1BB_Apt3T expression in D24-AptT-infected cells (MDA-BoM-1833) normalized to GAPDH (n = 3; mean ± SEM; paired t test). Fold changes are calculated relative to expression levels at 16 h. **B)** Correlation between viral mRNA gene expression (fiber) and 4-1BB_Apt3T in D24-AptT-infected cells (MDA-BoM-1833) at 16 h (blue) and 72 h (orange) after infection (Pearson). **C)** Fold change in fiber expression at 72 h relative to 16 h in MDA-BoM-1833 cells infected with D24-RGD, D24-ScrT or D24-AptT normalized to GAPDH (n = 3; mean ± SEM; one-way ANOVA, p = 0.56). **D and E)** Pull-down of 4-1BB_Apt3T from cell pellets (D) and supernatants (E) of MDA-BoM-1833 cultures infected with Delta-24-AptT (MOI 25) for 72 hours. Recombinant 4-1BB protein or a control protein (hIgG) was used for pull-down assays. Values correspond to the relative abundance of 4-1BB_Apt3T aptamer normalized to the input sample (2^-ΔΔCt). **F)** Histograms displaying the expression of the T cell activation marker CD25 in lymphocytes treated with bulk RNA obtained from mock or infected cells. The values indicate the percentage of positive cells in the test condition (red). The color legend is shown in each plot header.

Pull-down assays using recombinant 4-1BB protein demonstrated target-specific recovery of the 4-1BB_Apt3T aptamer from the bulk RNA of Delta-24-AptT-infected cells (**Fig 2D**). Importantly, the aptamer was also recovered from the extracellular media 72 hours after the oncolytic infection, as confirmed by pull-down in the supernatant fraction (**Fig 2E**). To evaluate the ability of Delta-24-AptT to stimulate T-cell responses, CD3-activated T cells were treated with bulk RNA from Delta-24-AptT, Delta-24-ScrT-, or mock-infected cells. RNA from Delta-24-AptT-infected cells induced an upregulation of the T-cell activation marker CD25 (65.5 %), compared to RNA from Delta-24-ScrT-infected cells (32.0 %), which was indistinguishable from mock-infected cells (**Fig 2F**). These findings confirm that Delta-24-AptT effectively infects cells and produces a functional 4-1BB_Apt3T aptamer that binds 4-1BB and enhances T-cell responses. Furthermore, our results corroborate that the addition of scaffold sequences (Tornado and linker) does not compromise the activity of the aptamer and validate Delta-24-RGD as a vector for the in situ production of functional aptamers.

In vitro, no significant differences in viral genome levels were observed among Delta-24-RGD, Delta-24-ScrT, and Delta-24-AptT at the early stage of infection (16 h), indicating that the aptamer construct does not impact virus infectivity (**Fig 3A**). Similarly, viral genome levels at 72 h post-infection (**Fig 3A**) showed no significant differences, with mean amplification rates of 7665-, 6913- and 3852-fold for Delta-24-RGD, Delta-24-ScrT and Delta-24-AptT, respectively, between 16 and 72 h (**Fig 3B**). These results confirmed the absence of significant differences in the production of infectious viral particles **(Fig. 3C)**, further validating that the transgenes do not impair viral replication. In vivo, quantification of viral genomes in MDA-BoM-1833 subcutaneous tumors demonstrated the sustained presence of Delta-24-AptT within the tumor mass **(Fig. 3D)** with a steady average of 2,000–3,000 viral genomes per ng of tumor DNA during the first two weeks after injection.

**Figure 3.**
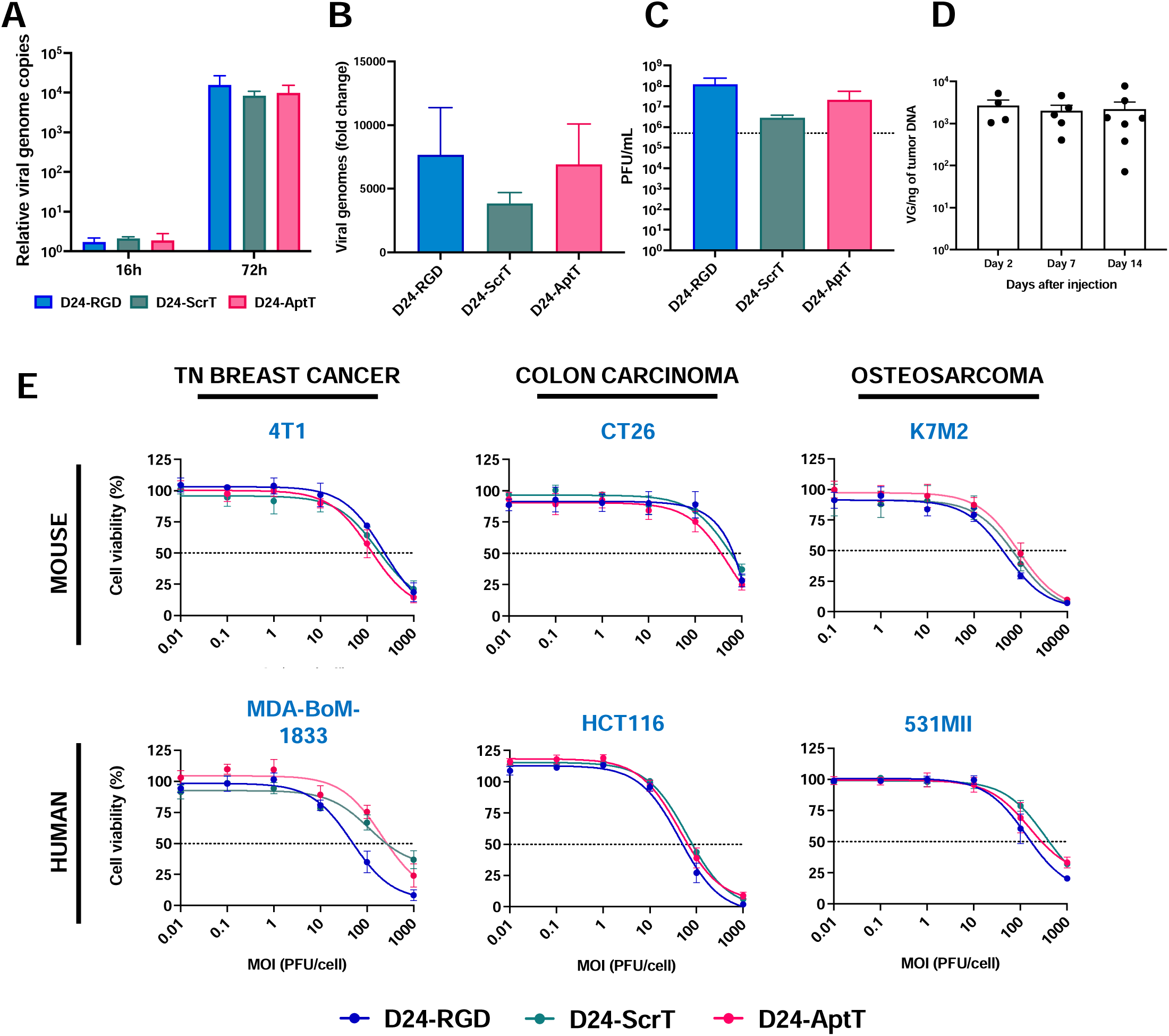
Preserved replicative and oncolytic properties of the modified virus. **A)** Viral genome copies normalized by total genomic DNA (2^-ΔCt) at in MDA-BoM-1833 cells at 16 h and 72 h after infection with Delta-24-RGD, Delta-24-ScrT or Delta-24-AptT (n = 3, mean ± SEM, two-way ANOVA, p = 0.75). **B)** Fold change in viral genomes at 72 h relative to 16 h (n = 3; mean ± SEM; one-way ANOVA , p = 0.63). **C)** Virus infectious titers (PFU) at 72 h after infection with the oncolytic viruses (n = 3; mean ± SEM; one-way ANOVA, p = 0.18). Dashed line indicates the virus input. **D)** Delta-24-AptT viral genomes (VG) per total tumor DNA at different time points after injection (n = 4-7; mean ± SEM; one-way ANOVA, p = 0.91). **E)** Cell viability in a panel of mouse and human tumor cell culture models three days after infection with increasing doses of each oncolytic virus, expressed as percentage relative to mock-infected cultures (n = 3; mean ± SEM). See also Supplemental Table S3.

Finally, assessing the oncolytic potency of Delta-24-AptT and Delta-24-ScrT compared to the parental Delta-24-RGD is essential to ensure that the therapeutic modifications do not compromise efficacy. Using a panel of mouse and human cell lines from different lineages, we evaluated the performance of the viruses across diverse tumor models (**Supplemental Table S3, Fig 3E**). In general, IC□□ values were higher in mouse models compared to their human counterparts across all three viruses, as expected due to the limited replication of human adenoviruses in most rodent cells^26^. Despite this, no significant differences in IC□□ values were observed among the three viruses in mouse models, with the cell model being the primary factor determining sensitivity. In human cell lines, Delta-24-AptT and Delta-24-ScrT exhibited potency comparable to that of Delta-24-RGD in most models, including HCT116 (colon carcinoma) and 531MII (osteosarcoma). Surprisingly, in MDA-BoM-1833 (breast carcinoma), Delta-24-AptT (IC□□ = 268.9, CI: 142.0–467.6) and Delta-24-ScrT (IC□□ = 258.6, CI: 117.8–823.4) showed reduced potency compared to Delta-24-RGD (IC□□ = 50.2, CI: 31.3–78.2). This cell-specific observation does not compromise the overall conclusion that Delta-24-AptT retains robust oncolytic activity across diverse models.

### Intratumor injection of Delta-24-AptT in vivo results in tumor growth control mediated by 4-1BB

Building upon our in vitro studies demonstrating the ability of Delta-24-AptT to produce functional 4-1BB_Apt3T aptamers and induce a cytopathic effect, we next evaluated its therapeutic efficacy in vivo. Mice bearing subcutaneous 4T1 tumors were treated with a single intratumoral injection of PBS (mock), Delta-24-ScrT (control virus), or Delta-24-AptT. Delta-24-AptT significantly reduced tumor growth (**Fig 4A and Supplemental Fig S5A**) compared to PBS (p = 0.0003) and Delta-24-ScrT (p = 0.0004). Individual tumor growth patterns further highlighted a consistent tumor-suppressive response in Apt-treated mice (**Fig 4B**), whereas tumor growth in Delta-24-ScrT was indistinguishable from that of the mock-treated group. Endpoint histological characterization of TILs revealed a higher total CD3+ T-cell infiltration in the aptamer-encoding virus (**Fig 4C and 4D**) and a significant increase in CD8+ T cells compared to mock (>3-fold; p = 0.001) and the control virus (>2-fold; p = 0.006), suggesting an enhanced cytotoxic immune response.

**Figure 4.**
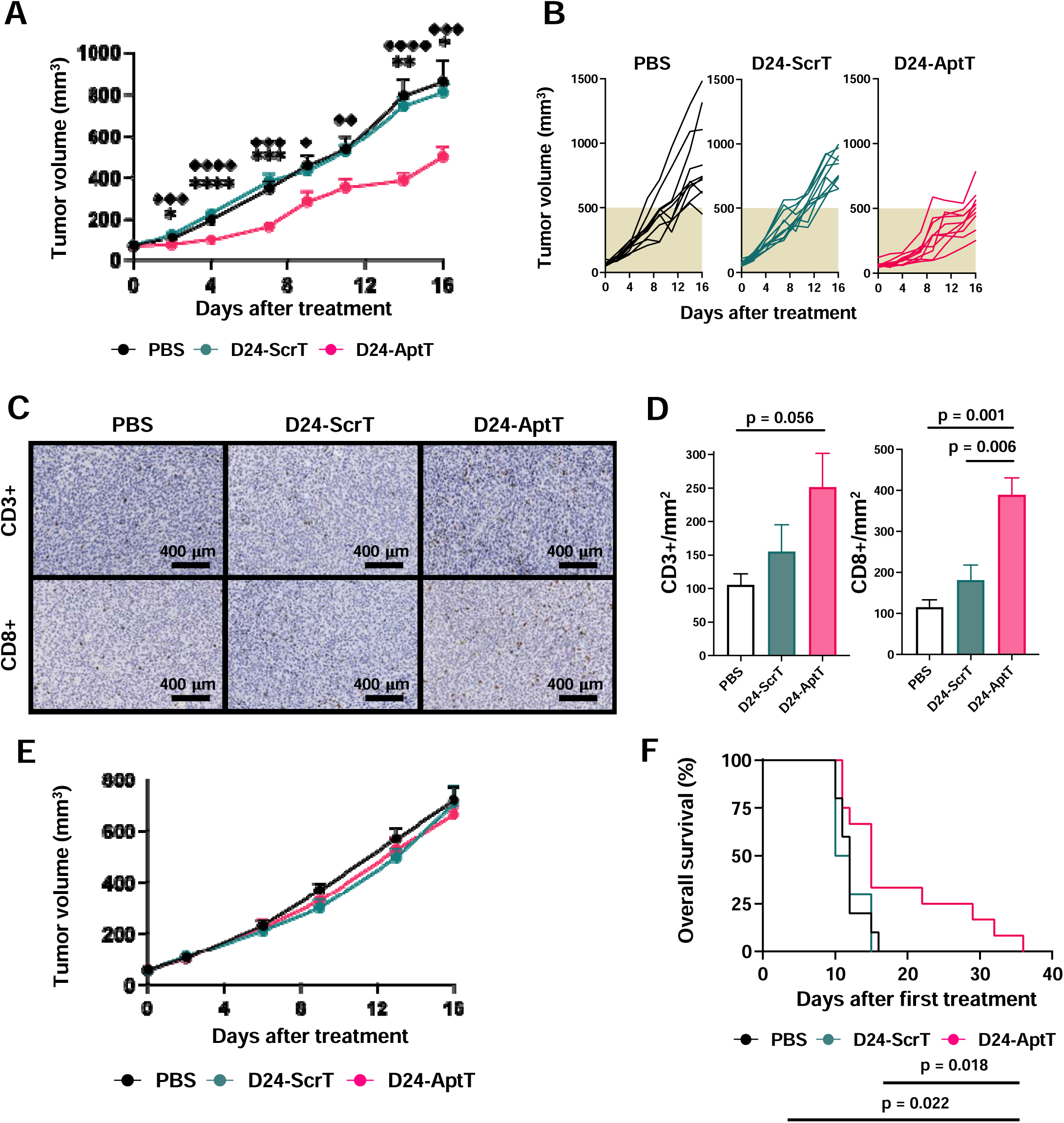
Therapeutic benefit of Delta-24-Apt3T in mouse in vivo tumor models. **A)** 4T1 tumor volume over time after being treated with the oncolytic viruses or PBS (n = 10; mean ± SEM, repeated two-way ANOVA, p < 0.0001). □ and □ indicate the statistical differences between Delta-24-AptT and Delta-24-ScrT or PBS, respectively (* p < 0.05, ** p < 0.01, *** p < 0.001, **** p < 0.0001). **B)** Spider plot displaying 4T1 tumor growth per animal. **C)** Representative images showing CD3 and CD8 staining in 4T1 tumors. **D)** Quantification of CD3+ and CD8+ cells in FFPE 4T1 tumors (n = 4 for PBS and D24-ScrT and 3 for D24-AptT; mean ± SEM, one-way ANOVA, p = 0.065 and p = 0.0011 for CD3 and CD8, respectively). **E)** 4T1 tumor growth in 4-1BB KO mice (n =6-8; mean ± SEM, repeated two-way ANOVA, p = 0.38). **F)** Kaplan-Meier curve showing overall survival of mice bearing intratibial osteosarcoma tumors after receiving two intratumoral injections (day 0 and day 8) of the oncolytic viruses or PBS (n = 10 for PBS and D24-ScrT, n = 12 for D24-AptT; Log rank test, p = 0.02). See also Supplemental Figure S5.

The effect of Delta-24-AptT on tumor growth control was abrogated entirely in a 4-1BB knockout (KO) background model (BALB/c CD137KO; C.Cg-Tnfrsf9^tm1Byk^), as shown in the complete overlap of tumor size charts in PBS, Delta-24-ScrT, and Delta-24-AptT (**Fig 4E**), thus confirming that the Delta-24-AptT mechanism of action relies on the presence of 4-1BB in the tumor microenvironment. Importantly, no significant differences were observed between Delta-24-AptT and Delta-24-ACT (**Supplemental Fig S5B**), a modified Delta-24-RGD expressing the 4-1BBL previously validated in various solid tumor models^5,27,28^.

### Delta-24-AptT provides measurable antitumor activity and survival benefit across tumor models

To further reinforce the translational relevance of Delta-24-AptT, we tested its therapeutic effect in additional in vivo tumor models. In CT26 subcutaneous tumors, Delta□24□AptT significantly reduced tumor growth compared with PBS, with a homogeneous response across animals **(Supplemental Fig. S5C and D).**

Finally, we tested the efficacy of the virus in an orthotopic intratibial osteosarcoma using the aggressive K7M2 model that has a high penetrance in developing spontaneous lung metastases, the most critical problem in osteosarcoma. The therapeutic advantage of Delta-24-AptT becomes clear analyzing the survival (p = 0.02). While PBS- and Delta-24-ScrT-treated groups had median overall survival of 12 and 11 days, respectively, Delta-24-AptT significantly extended survival (p = 0.02 compared to PBS and control virus) to a median of 15 days. Furthermore, 25% of Delta-24-AptT-treated animals survived beyond day 22, and 17% exceeded 30 days (**Fig 4F**).

### Delta-24-AptT drives a cytotoxic immune response profile in the tumor microenvironment

To better dissect the tumor microenvironment remodeling exerted by Delta-24-AptT, TILs were isolated from subcutaneous 4T1 10 days after treatment and analyzed by flow cytometry (Fig 5) following the gating strategy described in **Supplemental Fig S6A**. Delta-24-AptT treatment led to an overall increased immune cell infiltration compared to both mock- and Delta-24-ScrT-treated groups (**Fig 5A**). Specifically, Delta-24-ApT increased NK cell density relative to PBS and the control virus. While both viruses increased T-cell infiltration, only Delta-24-AptT showed a significant difference compared to PBS, suggesting a more robust effect than the control virus. An in-depth breakdown of T-cell subsets showed comparable levels of conventional CD4+ T cells between Delta-24-ScrT and Delta-24-AptT, with the latter showing a modest enrichment in CD8+ T cells. Tregs were also slightly elevated in Delta-24-AptT-treated tumors, but the strong impact on CD4 and CD8 T cells resulted in an overall Treg contribution to the T-cell pool (**Supplemental Fig S6B and S6C**). T-cell proportions also indicate a tip balance to cytotoxic T-cell responses in Delta-24-AptT.

**Figure 5.**
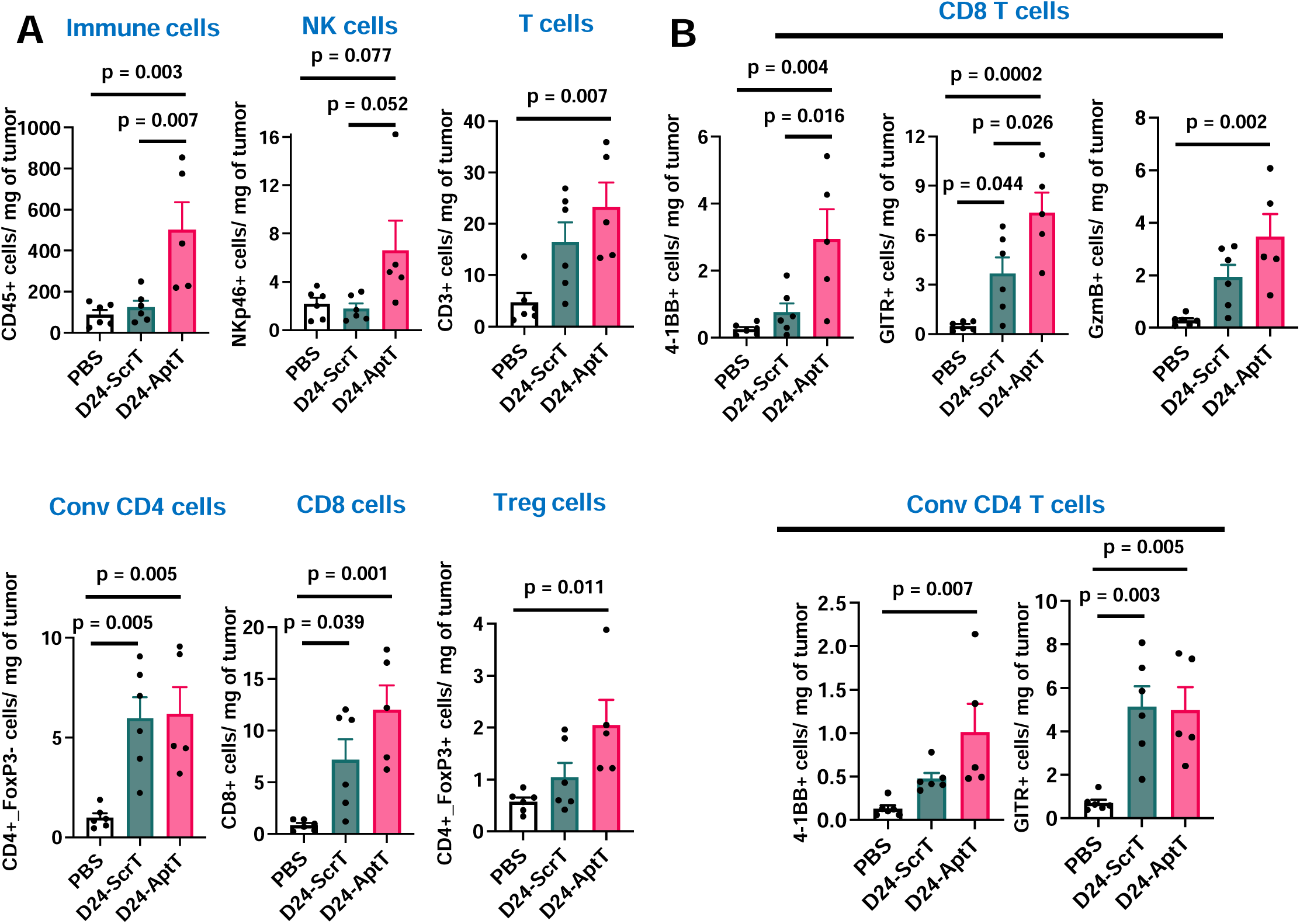
Delta-24-Apt3T induces proinflammatory remodeling of the tumor immune microenvironment. **A)** Immune cell profile of the lymphoid compartment in 4T1 tumors 10 days after treatment with Delta-24-AptT, Delta-24-ScrT or PBS. **B)** T cell counts expressing activation markers 4-1BB, GITR and GmzB (n = 5/6; mean ± SEM; one-way ANOVA). See also Supplemental Figure S6.

Focusing on T-cell activation and functional status, Delta-24-AptT demonstrated a distinct ability to enhance key activation markers compared with Delta-24-ScrT. Delta-24-AptT significantly increased the number of 4-1BB- and GITR-expressing CD8+ and conventional CD4+ T cells compared to PBS, and was also higher than the control virus in CD8 cells (**Fig 5B**). A pronounced functional activation was observed in the CD8+ subset, as evidenced by a significant enrichment of cells expressing the cytotoxic effector granzyme B in Delta-24-AptT-treated tumors.

Together, these results demonstrate that while both Delta-24-ScrT and Delta-24-AptT recruit immune cells into the tumor microenvironment, Delta-24-AptT uniquely enhances their activation and functional capacity. The significant enrichment of CD137+ and GITR+ T cells, particularly among CD8+ T cells, underscores the ability of the virus encoding 4-1BB aptamers to drive robust cytotoxic immune responses.

## DISCUSSION

Oncolytic virotherapy is an emergent class of immunotherapy^29^ that achieves tumor responses through direct oncolysis and induction of antitumor immunity. To date, three different oncolytic viruses have been approved by the regulatory agencies for the treatment of solid tumors^30–32^. To enhance their efficacy, oncolytic viruses can be modified to encode therapeutic transgenes for the delivery of immunostimulatory molecules in the TME, thus potentiating their immunotherapeutic arm, while avoiding the risk of systemic off-tumor toxicities.

Engineering oncolytic viruses to encode therapeutic monoclonal antibodies is a strategy that enables targeting any desirable druggable protein expressed in the TME. However, due to size constraints, this strategy is limited to viruses with very large packaging capacity^33^. Miniaturized antibody versions, such as single-domain antibodies (sdAbs), are small enough to be packed in viruses with smaller carrying capacity, like adenoviruses. However, the development of therapeutic nanobodies is expensive and time consuming since it involves the immunization of camelids with the desired target, thus complicating the drug discovery pipeline. RNA aptamers, on the other hand, are also small molecules and their generation do not require complex immunization protocols, thus representing an accessible alternative option. The small size of the aptamer cassette and the polymerase III promoter required for its transcription might even allow the use of a tandem of different immunostimulatory aptamers integrated within the same oncolytic virus. Furthermore, RNA aptamers do not encode any exogenous protein or peptide, thus displaying lower immunogenicity and avoiding the potential generation of neutralizing antibodies that limit the efficacy of the therapeutic molecule over time.

In this work we present a novel approach using oncolytic viruses as a delivery platform for immunotherapeutic aptamers targeting the TME. We have generated a prototype oncolytic adenovirus Delta-24-AptT expressing a 4-1BB agonist RNA aptamer and tested the feasibility of the strategy in various in vitro and in vivo tumor models. The engineering process addressed key challenges, including designing a dimerization element, and incorporating circularization sequences to protect the aptamer from exoribonuclease attack in the extracellular environment. While a previous study has utilized adenoviral vectors for in situ transcription of a therapeutic RNA aptamer that targets an endogenous protein^34^, our work is the first to use replicative viruses, leveraging virus-mediated oncolysis to deliver bioactive RNA aptamers to the TME, opening new avenues for virus-based cancer immunotherapy. The aim of our work was to serve as a proof of concept that an endogenous-expressed non-coding RNA molecule (aptamer) delivered by an oncolytic virus can exert receptor-specific extracellular immunostimulatory activity. This technology could be eventually applied to a new generation of oncolytic viruses armed with a palette of immunostimulatory RNA aptamers.

Overall, our findings demonstrate that the inclusion of expression cassettes of RNA aptamers constructs does not impair the infectivity, replication, or oncolytic efficacy of the virus across six different tumor models. We also show that the aptamer produced by Delta-24-AptT is fully functional and mediates antitumor effects across different in vivo models through a mechanism dependent on 4-1BB activation in the TME. The aptamer-expressing virus induces a pro-inflammatory shift, increasing the infiltration of NK and CD8+ T cells expressing activation and effector markers compared to the control virus. This immune remodeling aligns with the expected outcomes of 4-1BB pathway stimulation previously reported in studies using 4-1BBL-expressing oncolytic viruses^5,35^. Importantly, no significant differences were observed between the Delta-24-RGD derivatives expressing the 4-1BB agonistic aptamer and 4-1BBL. However, a limitation of this study is the lack of virus replication in murine cells, which restricts evaluation of the dual mechanism of action (oncolysis and aptamer function) in a single model. In a replication-permissive environment, such as human cells, enhanced effects are anticipated due to increased aptamer expression, accelerated oncolysis, and prolonged viral persistence. Future studies are needed to validate this hypothesis.

In summary, our results establish Delta-24-AptT as a versatile therapeutic platform that combines robust oncolytic activity with the ability to deliver therapeutic aptamers. This approach provides a foundation for the development of advanced oncolytic viruses tailored to target specific pathways and enhance antitumor immunity.

## DATA AVAILABILITY

The data that support the findings of this study are available within the paper or supplemental information, or source data file or available from the corresponding author upon request.

## ACKNOWLEDGEMENTS

The performed work was supported through a Plan de colaboración Internacional (PCI2021-122084-2B) Spanish Ministry of Science and Innovation and (SL). GRANATE project funded by the Government of Navarre in the frame of "Proyectos Estratégicos de I+D 2022-2025, Reto GEMA (001-1411-2022-000066)" (MMA, APG, SL). Instituto de Salud Carlos III y Fondos Feder (CP24CIII/00003 MGM, PI21/00940 APG, PI23/01155 MMA, PI23/01120 FP, PI18/00164 APG “A way to make Europe”); Fundación El sueño de Vicky; Asociación Pablo Ugarte-Fuerza Julen, Fundación ADEY, Fundación ACS, (APG and MMA), Fundación Hay que tomarse la vida con tumor; Fundación + Investigación + Vida (La Guareña). Red Española de Terapias Avanzadas TERAV ISCIII (RD24/0014/0004; Financiado por la Unión Europea-Next Generation EU. Plan de Recuperación Transformación y Resiliencia). Estratégicos Asociación Española Contra el Cáncer (AECC PRYES235096PAST FP). This project also received funding from the European Research Council (ERC) under the European Uniońs Horizon 2020 Research and Innovation Programme (817884 ViroPedTher to MMA).

## AUTHOR CONTRIBUTIONS

ACTC, FP, MMA, and MGM conceptualized the project. ACTC, VL, IAM, DdlN, SL, MGH, MZ, HV, and MGM conducted experiments and data acquisition. ACTC and MGM analyzed data and drafted the manuscript. ACTC, VL, IAM, DdlN, SL, MGH, MZ, APG, HV, JF, CGM, IM, FP, MMA, and MGM reviewed and edited the manuscript. FP, MMA, and MGM supervised the study. MMA managed funding, project oversight, and reporting.

## ETHICAL APPROVAL

The Ethical Committee of the University of Navarra (CEEA) approved all animal protocols performed in this study (E49-22(027-22E1, 015-23, 094-23). In vivo experiments were performed at the veterinary facilities of CIMA following the appropriate regulatory and ethical guidelines for experimental animal care.

## REFERENCES

1 Jiang H, Clise-Dwyer K, Ruisaard KE, Fan X, Tian W, Gumin J et al. Delta-24-RGD oncolytic adenovirus elicits anti-glioma immunity in an immunocompetent mouse model. PLoS One 2014; 9: e97407.

2 Lang FF, Conrad C, Gomez-Manzano C, Alfred Yung WK, Sawaya R, Weinberg JS et al. Phase I study of DNX-2401 (delta-24-RGD) oncolytic adenovirus: replication and immunotherapeutic effects in recurrent malignant glioma. Journal of Clinical Oncology 2018; 36: 1419–1427.

3 Jiang H, Shin DH, Nguyen TT, Fueyo J, Fan X, Henry V et al. Localized Treatment with Oncolytic Adenovirus Delta-24-RGDOX Induces Systemic Immunity against Disseminated Subcutaneous and Intracranial Melanomas. Clinical Cancer Research 2019; 25: 6801–6814.

4 Rivera-Molina Y, Jiang H, Fueyo J, Nguyen T, Shin DH, Youssef G et al. GITRL-armed Delta-24-RGD oncolytic adenovirus prolongs survival and induces anti-glioma immune memory. Neurooncol Adv 2019; 1: 1–11.

5 Laspidea V, Puigdelloses M, Labiano S, Marrodán L, Garcia-Moure M, Zalacain M et al. Exploiting 4-1BB immune checkpoint to enhance the efficacy of oncolytic virotherapy for diffuse intrinsic pontine gliomas. JCI Insight 2022; 7. doi:10.1172/JCI.INSIGHT.154812.

6 Musher BL, Rowinsky EK, Smaglo BG, Abidi W, Othman M, Patel K et al. LOAd703, an oncolytic virus-based immunostimulatory gene therapy, combined with chemotherapy for unresectable or metastatic pancreatic cancer (LOKON001): results from arm 1 of a non-randomised, single-centre, phase 1/2 study. Lancet Oncol 2024; 25: 488–500.

7 Feodoroff M, Hamdan F, Antignani G, Feola S, Fusciello M, Russo S et al. Enhancing T-cell recruitment in renal cell carcinoma with cytokine-armed adenoviruses. Oncoimmunology 2024; 13. doi:10.1080/2162402X.2024.2407532.

8 Pérez-Larraya JG, Garcia-Moure M, Labiano S, Patiño-García A, Dobbs J, Gonzalez-Huarriz M et al. Oncolytic DNX-2401 Virus for Pediatric Diffuse Intrinsic Pontine Glioma. N Engl J Med 2022; 386: 2471–2481.

9 Bett AJ, Prevec L, Graham FL. Packaging capacity and stability of human adenovirus type 5 vectors. J Virol 1993; 67: 5911–5921.

10 Paul S, Konig MF, Pardoll DM, Bettegowda C, Papadopoulos N, Wright KM et al. Cancer therapy with antibodies. Nature Reviews Cancer 2024 24:6 2024; 24: 399–426.

11 Suntharalingam G, Perry MR, Ward S, Brett SJ, Castello-Cortes A, Brunner MD et al. Cytokine Storm in a Phase 1 Trial of the Anti-CD28 Monoclonal Antibody TGN1412. New England Journal of Medicine 2006; 355: 1018–1028.

12 Chester C, Sanmamed MF, Wang J, Melero I. Immunotherapy targeting 4-1BB: mechanistic rationale, clinical results, and future strategies. Blood 2018; 131: 49–57.

13 De Martin E, Michot JM, Rosmorduc O, Guettier C, Samuel D. Liver toxicity as a limiting factor to the increasing use of immune checkpoint inhibitors. JHEP Reports 2020; 2. doi:10.1016/J.JHEPR.2020.100170/ASSET/D4A2A2FD-D540-43AE-B8EA-8D4590EA8258/MAIN.ASSETS/GR5.JPG.

14 Melero I, Castanon E, Alvarez M, Champiat S, Marabelle A. Intratumoural administration and tumour tissue targeting of cancer immunotherapies. Nature Reviews Clinical Oncology 2021 18:9 2021; 18: 558–576.

15 Shigdar S, Schrand B, Giangrande PH, de Franciscis V. Aptamers: Cutting edge of cancer therapies. Molecular Therapy 2021; 29: 2396–2411.

16 Ellington AD, Szostak JW. In vitro selection of RNA molecules that bind specific ligands. Nature 1990 346:6287 1990; 346: 818–822.

17 Kwon DSV& BS, *. 4-1BB (CD137), an inducible costimulatory receptor, as a specific target for cancer therapy. BMB Rep 2014; 47: 122–129.

18 Melero I, Shuford WW, Newby SA, Aruffo A, Ledbetter JA, Hellström KE et al. Monoclonal antibodies against the 4-1BB T-cell activation molecule eradicate established tumors. Nature Medicine 1997 3:6 1997; 3: 682–685.

19 Chester C, Sanmamed MF, Wang J, Melero I. Immunotherapy targeting 4-1BB: mechanistic rationale, clinical results, and future strategies. Blood 2018; 131: 49–57.

20 Watts TH, Yeung KKM, Yu T, Lee S, Eshraghisamani R. TNF/TNFR Superfamily Members in Costimulation of T Cell Responses—Revisited. Annu Rev Immunol 2025. doi:10.1146/ANNUREV-IMMUNOL-082423-040557.

21 McNamara JO, Kolonias D, Pastor F, Mittler RS, Chen L, Giangrande PH et al. Multivalent 4-1BB binding aptamers costimulate CD8+ T cells and inhibit tumor growth in mice. J Clin Invest 2008; 118: 376–86.

22 Melero I, Murillo O, Dubrot J, Hervás-Stubbs S, Perez-Gracia JL. Multi-layered action mechanisms of CD137 (4-1BB)-targeted immunotherapies. Trends Pharmacol Sci 2008; 29: 383–390.

23 Singh R, Kim YH, Lee SJ, Eom HS, Choi BK. 4-1BB immunotherapy: advances and hurdles. Experimental & Molecular Medicine 2023 56:1 2024; 56: 32–39.

24 Zapata JM, Perez-Chacon G, Carr-Baena P, Martinez-Forero I, Azpilikueta A, Otano I et al. CD137 (4-1BB) signalosome: Complexity is a matter of TRAFs. Front Immunol 2018; 9: 422997.

25 Litke JL, Jaffrey SR. Highly efficient expression of circular RNA aptamers in cells using autocatalytic transcripts. Nat Biotechnol 2019. doi:10.1038/s41587-019-0090-6.

26 Blair GE, Dixon SC, Griffiths SA, Zajdel ME. Restricted replication of human adenovirus type 5 in mouse cell lines. Virus Res 1989; 14: 339–346.

27 Puigdelloses M, Garcia-Moure M, Labiano S, Laspidea V, Gonzalez-Huarriz M, Zalacain M et al. CD137 and PD-L1 targeting with immunovirotherapy induces a potent and durable antitumor immune response in glioblastoma models. J Immunother Cancer 2021; 9: e002644.

28 Martinez-Velez N, Laspidea V, Zalacain M, Labiano S, García-Moure M, Puigdelloses M et al. Local Treatment of a Pediatric Osteosarcoma Model with a 4-1BBL Armed Oncolytic Adenovirus Results in an Antitumor Effect and Leads to Immune Memory. Mol Cancer Ther 2022; 21: 471–480.

29 Kaufman HL, Kohlhapp FJ, Zloza A. Oncolytic viruses: a new class of immunotherapy drugs. Nature Reviews Drug Discovery 2015 14:9 2015; 14: 642–662.

30 Garber K. China approves world’s first oncolytic virus therapy for cancer treatment. J Natl Cancer Inst 2006; 98: 298–300.

31 Pol J, Kroemer G, Galluzzi L. First oncolytic virus approved for melanoma immunotherapy. Oncoimmunology 2015; 5: e1115641.

32 Frampton JE. Teserpaturev/G47Δ: First Approval. BioDrugs 2022; 36: 667–672.

33 Xu B, Tian L, Chen J, Wang J, Ma R, Dong W et al. An oncolytic virus expressing a full-length antibody enhances antitumor innate immune response to glioblastoma. Nature Communications 2021 12:1 2021; 12: 1–17.

34 Mi J, Zhang X, Rabbani ZN, Liu Y, Reddy SK, Su Z et al. RNA aptamer-targeted inhibition of NF-κB suppresses non-small cell lung cancer resistance to doxorubicin. Molecular Therapy 2008; 16: 66–73.

35 Hinterberger M, Giessel R, Fiore G, Graebnitz F, Bathke B, Wennier S et al. Intratumoral virotherapy with 4-1BBL armed modified vaccinia Ankara eradicates solid tumors and promotes protective immune memory. J Immunother Cancer 2021; 9: e001586.

36 Alam KK, Chang JL, Burke DH. FASTAptamer: A bioinformatic toolkit for high-throughput sequence analysis of combinatorial selections. Mol Ther Nucleic Acids 2015; 4: e230.

37 Reuter JS, Mathews DH. RNAstructure: Software for RNA secondary structure prediction and analysis. BMC Bioinformatics 2010; 11: 1–9.

38 Popenda M, Szachniuk M, Antczak M, Purzycka KJ, Lukasiak P, Bartol N et al. Automated 3D structure composition for large RNAs. Nucleic Acids Res 2012; 40: e112–e112.

39 Sehnal D, Bittrich S, Deshpande M, Svobodová R, Berka K, Bazgier V et al. Mol* Viewer: modern web app for 3D visualization and analysis of large biomolecular structures. Nucleic Acids Res 2021; 49: W431–W437.

40 Cascallo M, Gros A, Bayo N, Serrano T, Capella G, Alemany R. Deletion of VAI and VAII RNA Genes in the Design of Oncolytic Adenoviruses. Hum Gene Ther 2006; 17: 929–940.

