## Supplementary material for "Replication-competent adenoviral platform for in situ production of immunotherapeutic RNA aptamers targeting 4-1BB"

### Slide 1
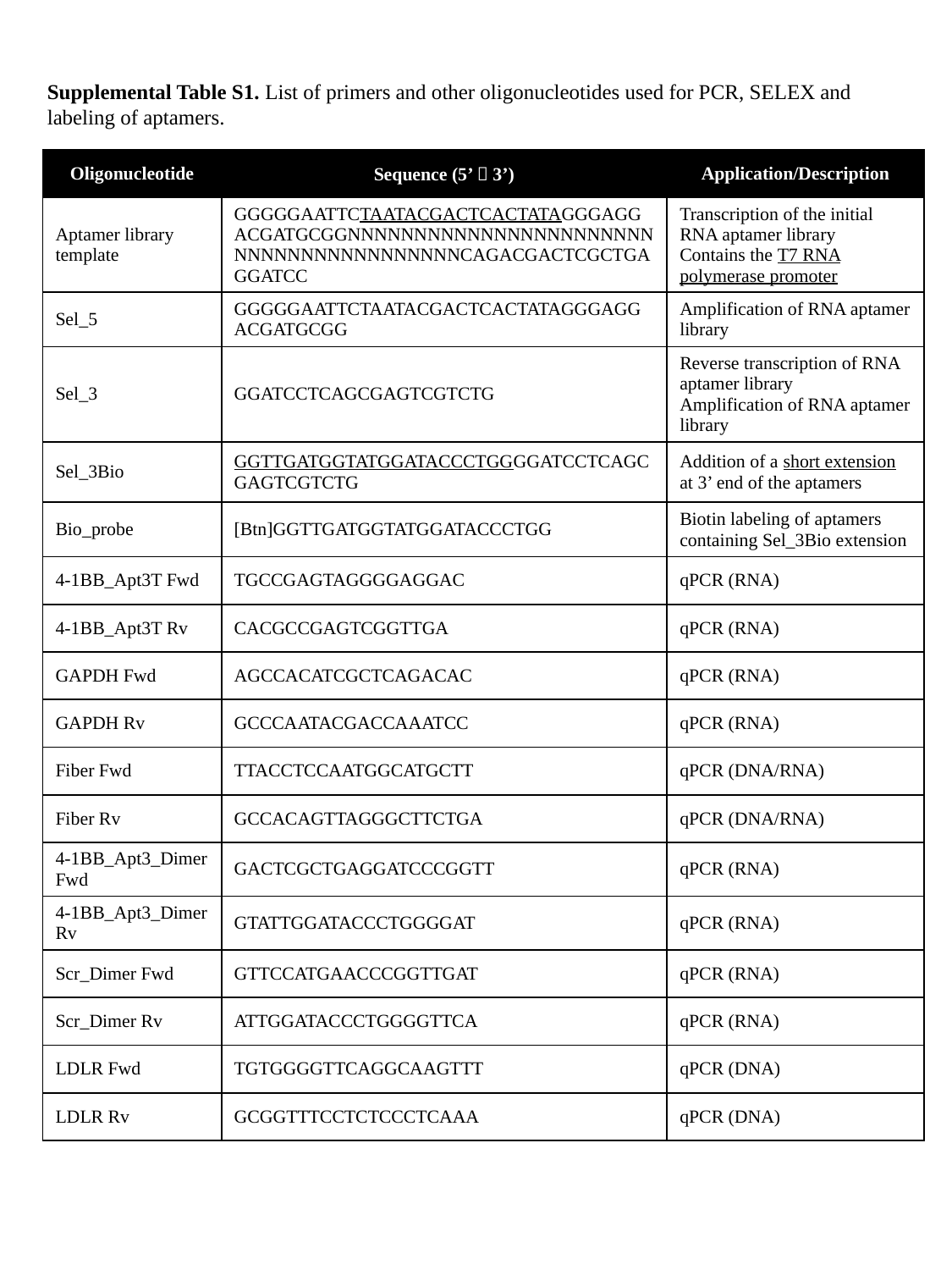

Supplemental Table S1. List of primers and other oligonucleotides used for PCR, SELEX and labeling of aptamers.
| Oligonucleotide | Sequence (5’ 🡪 3’) | Application/Description |
| --- | --- | --- |
| Aptamer library template | GGGGGAATTCTAATACGACTCACTATAGGGAGGACGATGCGGNNNNNNNNNNNNNNNNNNNNNNNNNNNNNNNNNNNNNNNNCAGACGACTCGCTGAGGATCC | Transcription of the initial RNA aptamer library Contains the T7 RNA polymerase promoter |
| Sel\_5 | GGGGGAATTCTAATACGACTCACTATAGGGAGGACGATGCGG | Amplification of RNA aptamer library |
| Sel\_3 | GGATCCTCAGCGAGTCGTCTG | Reverse transcription of RNA aptamer library Amplification of RNA aptamer library |
| Sel\_3Bio | GGTTGATGGTATGGATACCCTGGGGATCCTCAGCGAGTCGTCTG | Addition of a short extension at 3’ end of the aptamers |
| Bio\_probe | [Btn]GGTTGATGGTATGGATACCCTGG | Biotin labeling of aptamers containing Sel\_3Bio extension |
| 4-1BB\_Apt3T Fwd | TGCCGAGTAGGGGAGGAC | qPCR (RNA) |
| 4-1BB\_Apt3T Rv | CACGCCGAGTCGGTTGA | qPCR (RNA) |
| GAPDH Fwd | AGCCACATCGCTCAGACAC | qPCR (RNA) |
| GAPDH Rv | GCCCAATACGACCAAATCC | qPCR (RNA) |
| Fiber Fwd | TTACCTCCAATGGCATGCTT | qPCR (DNA/RNA) |
| Fiber Rv | GCCACAGTTAGGGCTTCTGA | qPCR (DNA/RNA) |
| 4-1BB\_Apt3\_Dimer Fwd | GACTCGCTGAGGATCCCGGTT | qPCR (RNA) |
| 4-1BB\_Apt3\_Dimer Rv | GTATTGGATACCCTGGGGAT | qPCR (RNA) |
| Scr\_Dimer Fwd | GTTCCATGAACCCGGTTGAT | qPCR (RNA) |
| Scr\_Dimer Rv | ATTGGATACCCTGGGGTTCA | qPCR (RNA) |
| LDLR Fwd | TGTGGGGTTCAGGCAAGTTT | qPCR (DNA) |
| LDLR Rv | GCGGTTTCCTCTCCCTCAAA | qPCR (DNA) |

### Slide 2
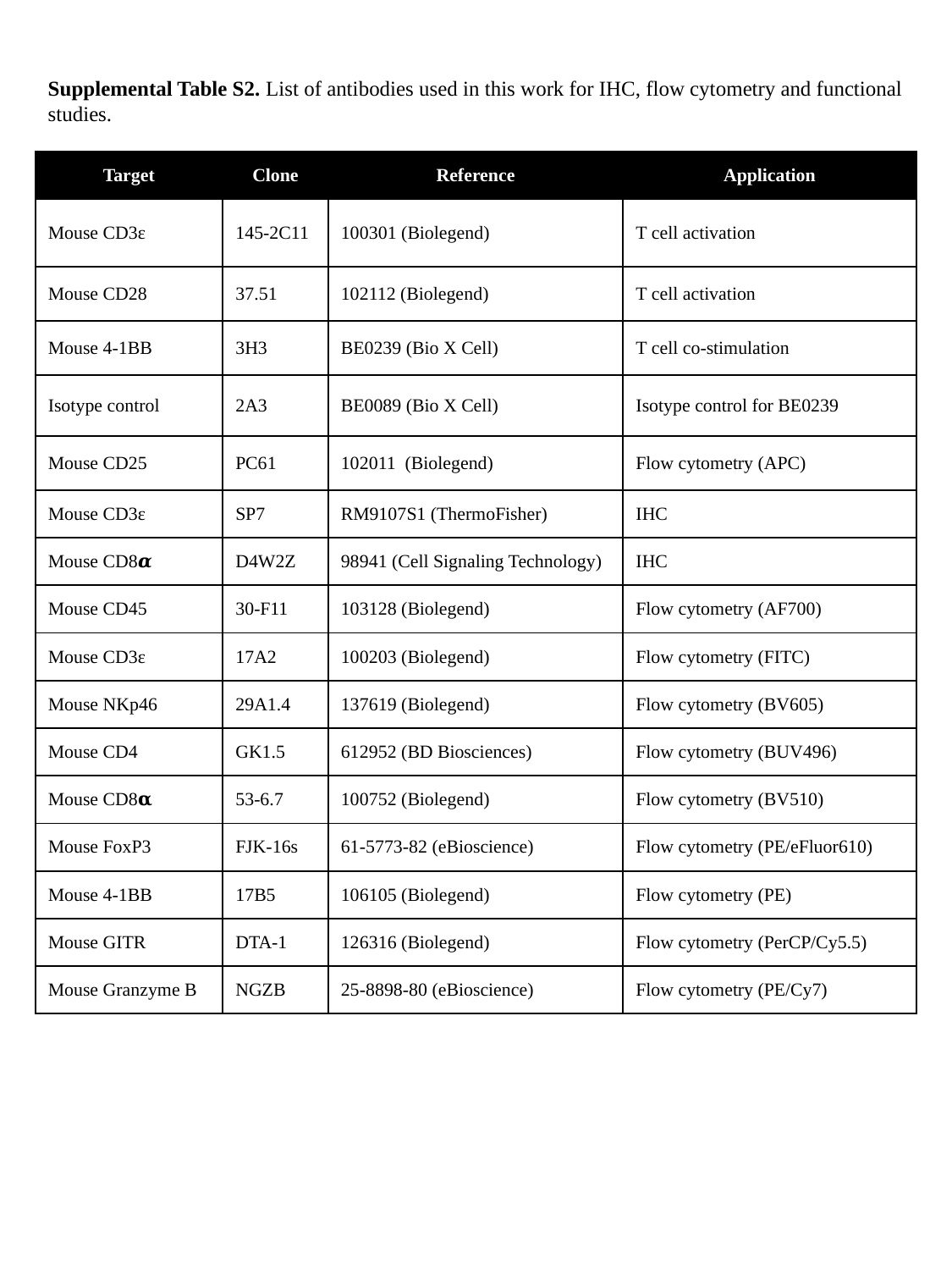

Supplemental Table S2. List of antibodies used in this work for IHC, flow cytometry and functional studies.
| Target | Clone | Reference | Application |
| --- | --- | --- | --- |
| Mouse CD3ɛ | 145-2C11 | 100301 (Biolegend) | T cell activation |
| Mouse CD28 | 37.51 | 102112 (Biolegend) | T cell activation |
| Mouse 4-1BB | 3H3 | BE0239 (Bio X Cell) | T cell co-stimulation |
| Isotype control | 2A3 | BE0089 (Bio X Cell) | Isotype control for BE0239 |
| Mouse CD25 | PC61 | 102011 (Biolegend) | Flow cytometry (APC) |
| Mouse CD3ɛ | SP7 | RM9107S1 (ThermoFisher) | IHC |
| Mouse CD8𝜶 | D4W2Z | 98941 (Cell Signaling Technology) | IHC |
| Mouse CD45 | 30-F11 | 103128 (Biolegend) | Flow cytometry (AF700) |
| Mouse CD3ɛ | 17A2 | 100203 (Biolegend) | Flow cytometry (FITC) |
| Mouse NKp46 | 29A1.4 | 137619 (Biolegend) | Flow cytometry (BV605) |
| Mouse CD4 | GK1.5 | 612952 (BD Biosciences) | Flow cytometry (BUV496) |
| Mouse CD8𝛂 | 53-6.7 | 100752 (Biolegend) | Flow cytometry (BV510) |
| Mouse FoxP3 | FJK-16s | 61-5773-82 (eBioscience) | Flow cytometry (PE/eFluor610) |
| Mouse 4-1BB | 17B5 | 106105 (Biolegend) | Flow cytometry (PE) |
| Mouse GITR | DTA-1 | 126316 (Biolegend) | Flow cytometry (PerCP/Cy5.5) |
| Mouse Granzyme B | NGZB | 25-8898-80 (eBioscience) | Flow cytometry (PE/Cy7) |

### Slide 3
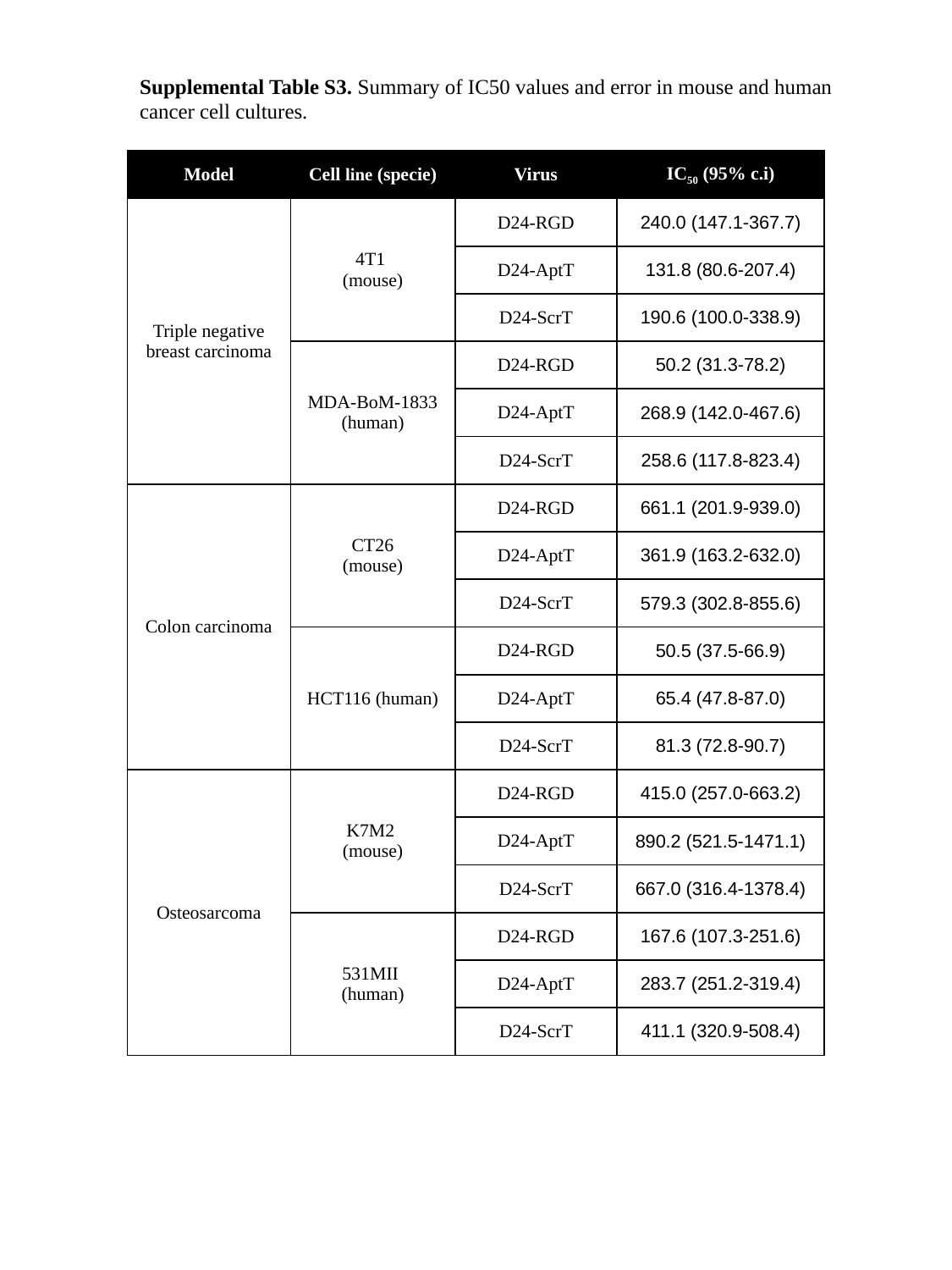

Supplemental Table S3. Summary of IC50 values and error in mouse and human cancer cell cultures.
| Model | Cell line (specie) | Virus | IC_50_ (95% c.i) |
| --- | --- | --- | --- |
| Triple negative breast carcinoma | 4T1 (mouse) | D24-RGD | 240.0 (147.1-367.7) |
| | | D24-AptT | 131.8 (80.6-207.4) |
| | | D24-ScrT | 190.6 (100.0-338.9) |
| | MDA-BoM-1833 (human) | D24-RGD | 50.2 (31.3-78.2) |
| | | D24-AptT | 268.9 (142.0-467.6) |
| | | D24-ScrT | 258.6 (117.8-823.4) |
| Colon carcinoma | CT26 (mouse) | D24-RGD | 661.1 (201.9-939.0) |
| | | D24-AptT | 361.9 (163.2-632.0) |
| | | D24-ScrT | 579.3 (302.8-855.6) |
| | HCT116 (human) | D24-RGD | 50.5 (37.5-66.9) |
| | | D24-AptT | 65.4 (47.8-87.0) |
| | | D24-ScrT | 81.3 (72.8-90.7) |
| Osteosarcoma | K7M2 (mouse) | D24-RGD | 415.0 (257.0-663.2) |
| | | D24-AptT | 890.2 (521.5-1471.1) |
| | | D24-ScrT | 667.0 (316.4-1378.4) |
| | 531MII (human) | D24-RGD | 167.6 (107.3-251.6) |
| | | D24-AptT | 283.7 (251.2-319.4) |
| | | D24-ScrT | 411.1 (320.9-508.4) |

### Slide 4
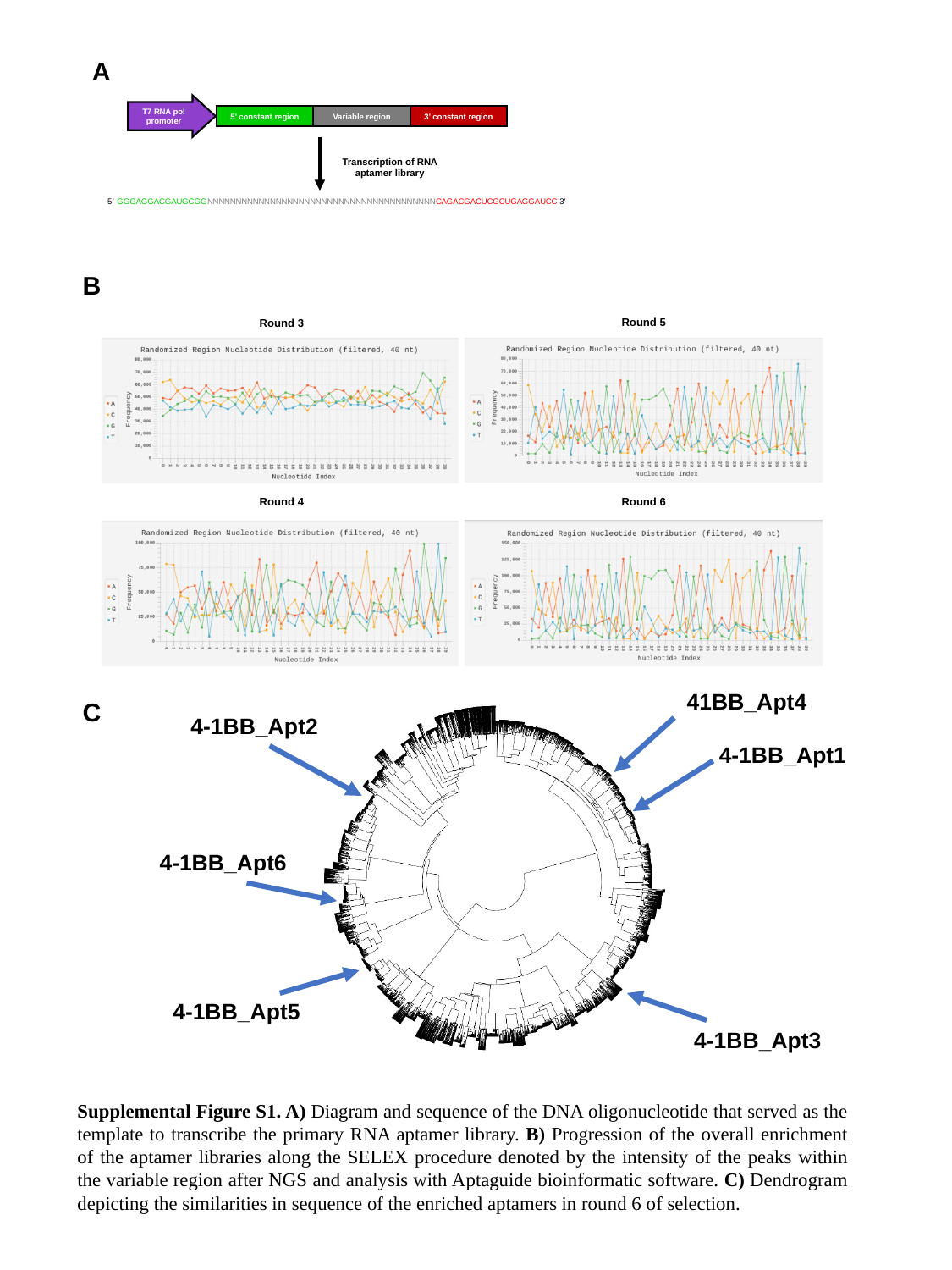

A
T7 RNA pol promoter
5’ constant region
Variable region
3’ constant region
Transcription of RNA aptamer library
5` GGGAGGACGAUGCGGNNNNNNNNNNNNNNNNNNNNNNNNNNNNNNNNNNNNNNNNCAGACGACUCGCUGAGGAUCC 3’
B
Round 5
Round 3
Round 4
Round 6
B
4-1BB_Apt1
4-1BB_Apt5
4-1BB_Apt3
4-1BB_Apt2
4-1BB_Apt6
41BB_Apt4
C
Supplemental Figure S1. A) Diagram and sequence of the DNA oligonucleotide that served as the template to transcribe the primary RNA aptamer library. B) Progression of the overall enrichment of the aptamer libraries along the SELEX procedure denoted by the intensity of the peaks within the variable region after NGS and analysis with Aptaguide bioinformatic software. C) Dendrogram depicting the similarities in sequence of the enriched aptamers in round 6 of selection.

### Slide 5
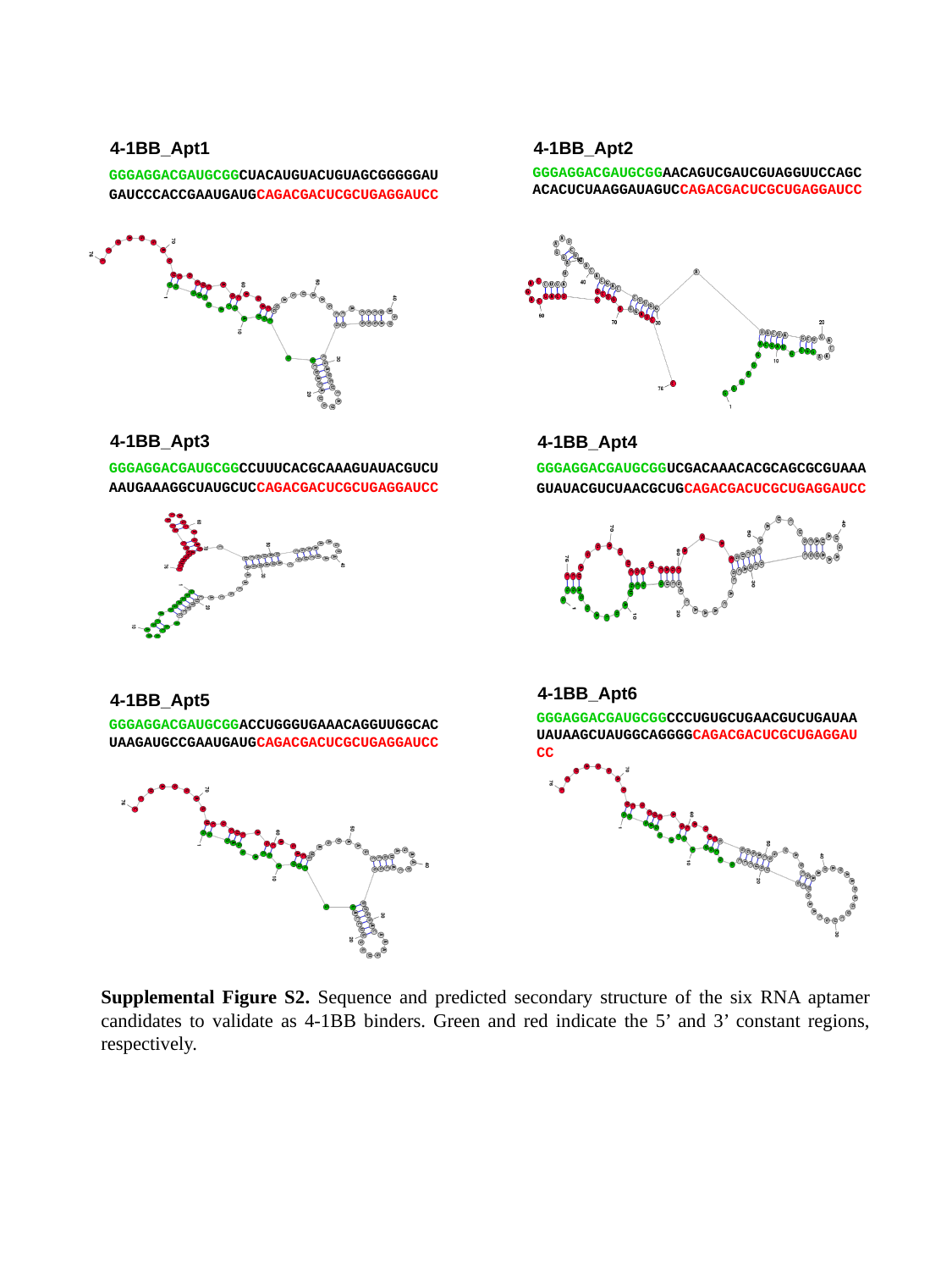

4-1BB_Apt1
4-1BB_Apt2
GGGAGGACGAUGCGGCUACAUGUACUGUAGCGGGGGAUGAUCCCACCGAAUGAUGCAGACGACUCGCUGAGGAUCC
GGGAGGACGAUGCGGAACAGUCGAUCGUAGGUUCCAGCACACUCUAAGGAUAGUCCAGACGACUCGCUGAGGAUCC
4-1BB_Apt3
4-1BB_Apt4
GGGAGGACGAUGCGGCCUUUCACGCAAAGUAUACGUCUAAUGAAAGGCUAUGCUCCAGACGACUCGCUGAGGAUCC
GGGAGGACGAUGCGGUCGACAAACACGCAGCGCGUAAAGUAUACGUCUAACGCUGCAGACGACUCGCUGAGGAUCC
4-1BB_Apt6
4-1BB_Apt5
GGGAGGACGAUGCGGCCCUGUGCUGAACGUCUGAUAAUAUAAGCUAUGGCAGGGGCAGACGACUCGCUGAGGAUCC
GGGAGGACGAUGCGGACCUGGGUGAAACAGGUUGGCACUAAGAUGCCGAAUGAUGCAGACGACUCGCUGAGGAUCC
Supplemental Figure S2. Sequence and predicted secondary structure of the six RNA aptamer candidates to validate as 4-1BB binders. Green and red indicate the 5’ and 3’ constant regions, respectively.

### Slide 6
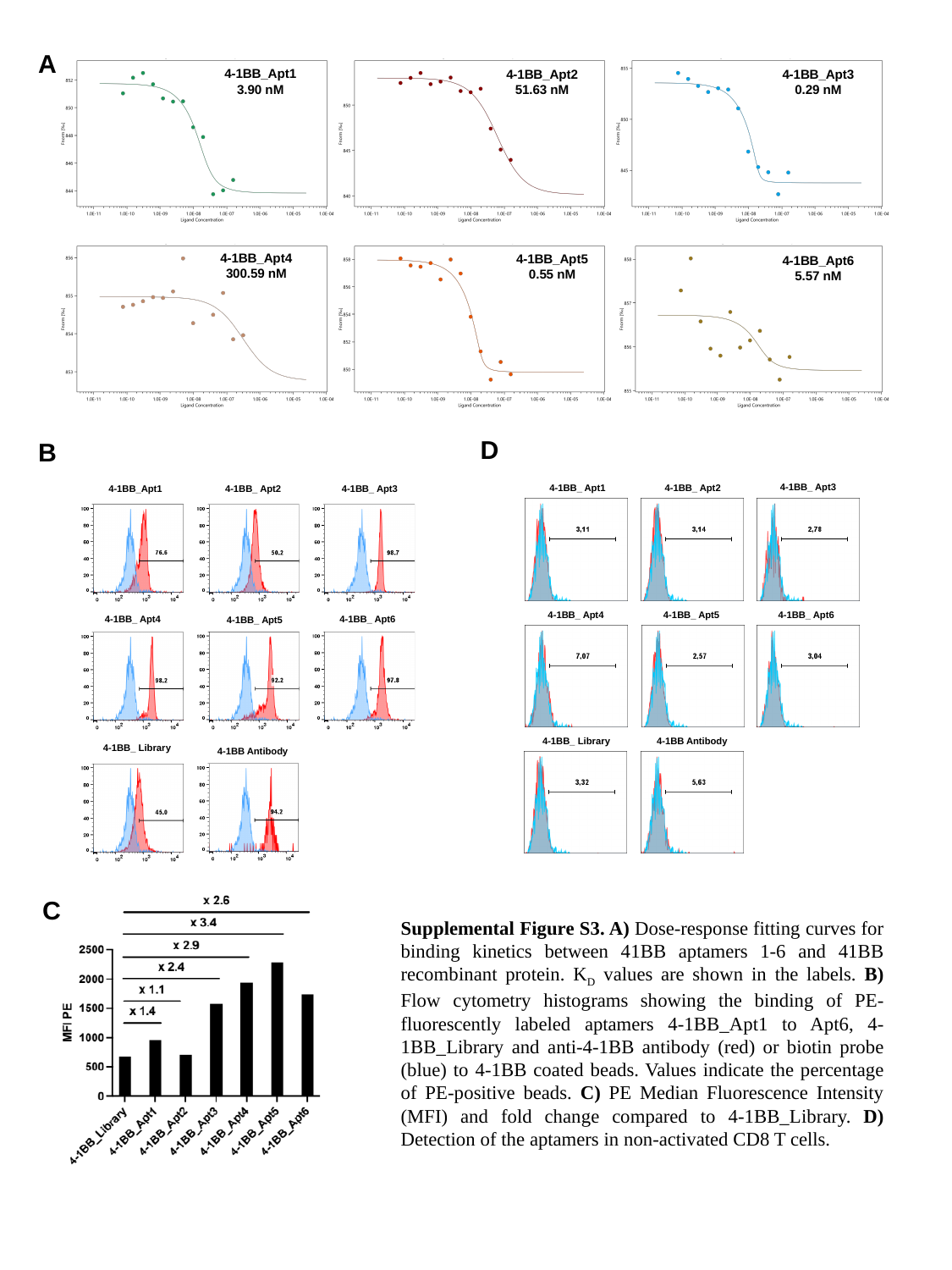

A
4-1BB_Apt1
3.90 nM
4-1BB_Apt2
51.63 nM
4-1BB_Apt3
0.29 nM
4-1BB_Apt4
300.59 nM
4-1BB_Apt5
0.55 nM
4-1BB_Apt6
5.57 nM
D
B
4-1BB_ Apt3
4-1BB_ Apt1
4-1BB_ Apt2
4-1BB_ Apt4
4-1BB_ Apt6
4-1BB_ Apt5
4-1BB_ Library
4-1BB Antibody
4-1BB_ Apt3
4-1BB_Apt1
4-1BB_ Apt2
4-1BB_ Apt4
4-1BB_ Apt6
4-1BB_ Apt5
4-1BB_ Library
4-1BB Antibody
C
Supplemental Figure S3. A) Dose-response fitting curves for binding kinetics between 41BB aptamers 1-6 and 41BB recombinant protein. K_D_ values are shown in the labels. B) Flow cytometry histograms showing the binding of PE-fluorescently labeled aptamers 4-1BB_Apt1 to Apt6, 4-1BB_Library and anti-4-1BB antibody (red) or biotin probe (blue) to 4-1BB coated beads. Values indicate the percentage of PE-positive beads. C) PE Median Fluorescence Intensity (MFI) and fold change compared to 4-1BB_Library. D) Detection of the aptamers in non-activated CD8 T cells.

### Slide 7
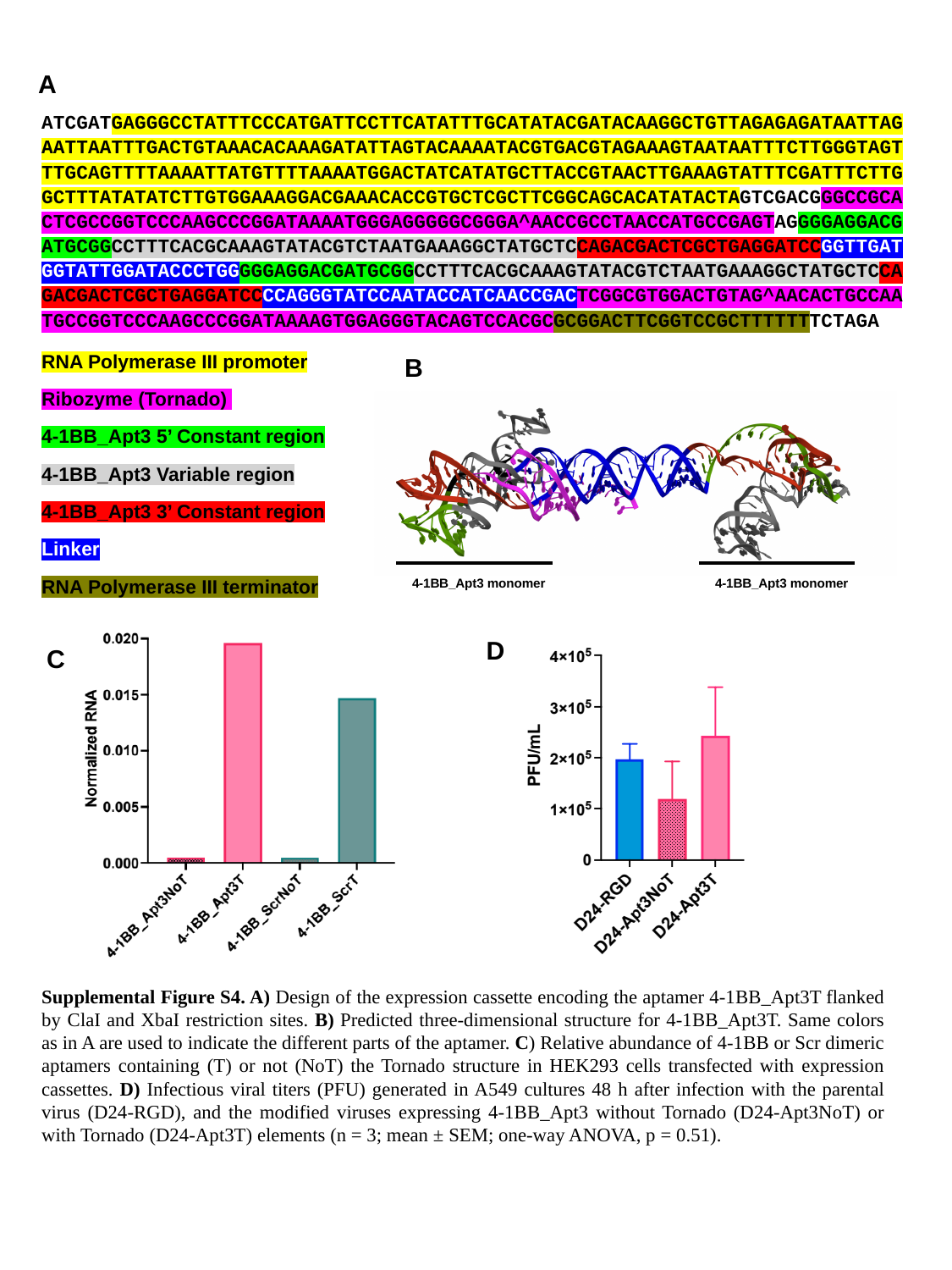

A
ATCGATGAGGGCCTATTTCCCATGATTCCTTCATATTTGCATATACGATACAAGGCTGTTAGAGAGATAATTAGAATTAATTTGACTGTAAACACAAAGATATTAGTACAAAATACGTGACGTAGAAAGTAATAATTTCTTGGGTAGTTTGCAGTTTTAAAATTATGTTTTAAAATGGACTATCATATGCTTACCGTAACTTGAAAGTATTTCGATTTCTTGGCTTTATATATCTTGTGGAAAGGACGAAACACCGTGCTCGCTTCGGCAGCACATATACTAGTCGACGGGCCGCACTCGCCGGTCCCAAGCCCGGATAAAATGGGAGGGGGCGGGA^AACCGCCTAACCATGCCGAGTAGGGGAGGACGATGCGGCCTTTCACGCAAAGTATACGTCTAATGAAAGGCTATGCTCCAGACGACTCGCTGAGGATCCGGTTGATGGTATTGGATACCCTGGGGGAGGACGATGCGGCCTTTCACGCAAAGTATACGTCTAATGAAAGGCTATGCTCCAGACGACTCGCTGAGGATCCCCAGGGTATCCAATACCATCAACCGACTCGGCGTGGACTGTAG^AACACTGCCAATGCCGGTCCCAAGCCCGGATAAAAGTGGAGGGTACAGTCCACGCGCGGACTTCGGTCCGCTTTTTTTCTAGA
RNA Polymerase III promoter
Ribozyme (Tornado)
4-1BB_Apt3 5’ Constant region
4-1BB_Apt3 Variable region
4-1BB_Apt3 3’ Constant region
Linker
RNA Polymerase III terminator
B
4-1BB_Apt3 monomer
4-1BB_Apt3 monomer
D
C
Supplemental Figure S4. A) Design of the expression cassette encoding the aptamer 4-1BB_Apt3T flanked by ClaI and XbaI restriction sites. B) Predicted three-dimensional structure for 4-1BB_Apt3T. Same colors as in A are used to indicate the different parts of the aptamer. C) Relative abundance of 4-1BB or Scr dimeric aptamers containing (T) or not (NoT) the Tornado structure in HEK293 cells transfected with expression cassettes. D) Infectious viral titers (PFU) generated in A549 cultures 48 h after infection with the parental virus (D24-RGD), and the modified viruses expressing 4-1BB_Apt3 without Tornado (D24-Apt3NoT) or with Tornado (D24-Apt3T) elements (n = 3; mean ± SEM; one-way ANOVA, p = 0.51).

### Slide 8
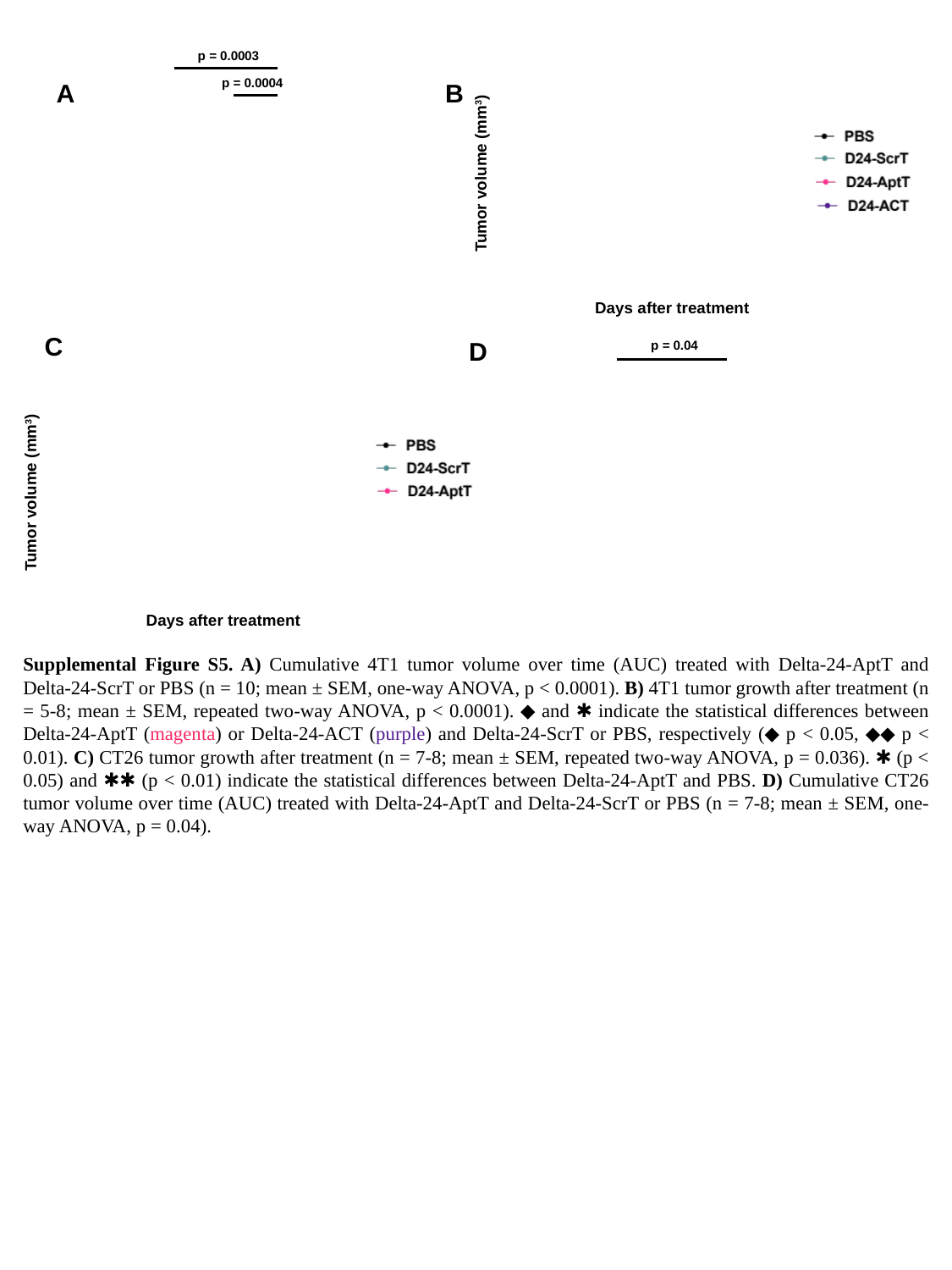

p = 0.0003
p = 0.0004
B
A
Tumor volume (mm^3^)
Days after treatment
C
D
p = 0.04
Tumor volume (mm^3^)
Days after treatment
Supplemental Figure S5. A) Cumulative 4T1 tumor volume over time (AUC) treated with Delta-24-AptT and Delta-24-ScrT or PBS (n = 10; mean ± SEM, one-way ANOVA, p < 0.0001). B) 4T1 tumor growth after treatment (n = 5-8; mean ± SEM, repeated two-way ANOVA, p < 0.0001). ◆ and ✱ indicate the statistical differences between Delta-24-AptT (magenta) or Delta-24-ACT (purple) and Delta-24-ScrT or PBS, respectively (◆ p < 0.05, ◆◆ p < 0.01). C) CT26 tumor growth after treatment (n = 7-8; mean ± SEM, repeated two-way ANOVA, p = 0.036). ✱ (p < 0.05) and ✱✱ (p < 0.01) indicate the statistical differences between Delta-24-AptT and PBS. D) Cumulative CT26 tumor volume over time (AUC) treated with Delta-24-AptT and Delta-24-ScrT or PBS (n = 7-8; mean ± SEM, one-way ANOVA, p = 0.04).

### Slide 9
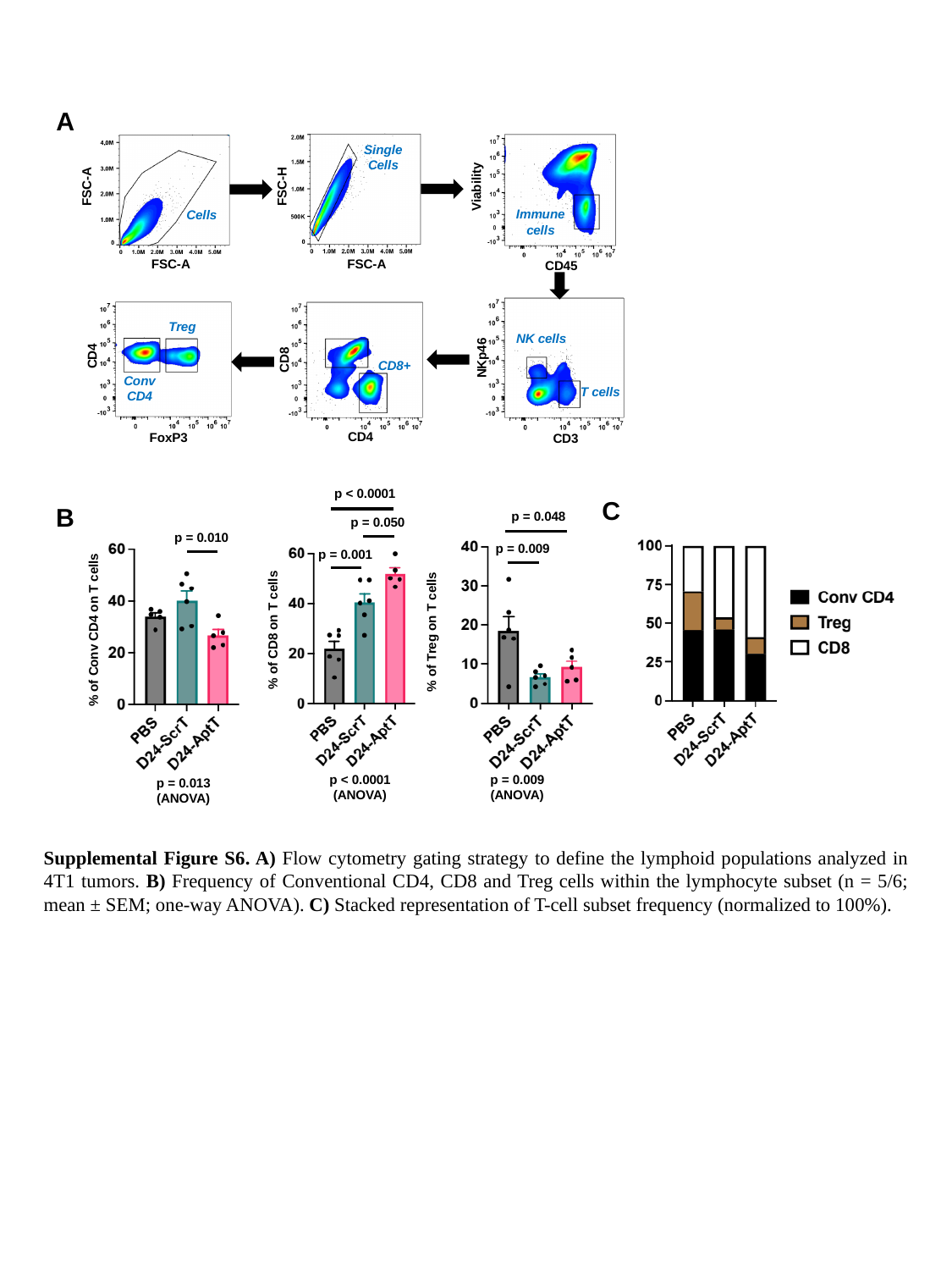

A
SingleCells
FSC-A
FSC-H
Viability
Cells
Immune cells
FSC-A
FSC-A
CD45
Treg
NK cells
CD4
NKp46
CD8
CD8+
Conv CD4
T cells
CD4
FoxP3
CD3
p < 0.0001
p = 0.048
p = 0.009
p = 0.050
p = 0.010
p = 0.001
% of Conv CD4 on T cells
% of CD8 on T cells
% of Treg on T cells
p < 0.0001
(ANOVA)
p = 0.009
(ANOVA)
p = 0.013
(ANOVA)
C
B
Supplemental Figure S6. A) Flow cytometry gating strategy to define the lymphoid populations analyzed in 4T1 tumors. B) Frequency of Conventional CD4, CD8 and Treg cells within the lymphocyte subset (n = 5/6; mean ± SEM; one-way ANOVA). C) Stacked representation of T-cell subset frequency (normalized to 100%).
